# 3D Bioprinted perfusable and vascularized breast tumor model for dynamic screening of chemotherapeutics and CAR-T cells

**DOI:** 10.1101/2022.03.15.484485

**Authors:** Madhuri Dey, Myong Hwan Kim, Momoka Nagamine, Mikail Dogan, Lina Kozhaya, Derya Unutmaz, Ibrahim T. Ozbolat

**Author notes:** Corresponding author: Ibrahim T. Ozbolat.

## Abstract

Despite substantial advancements in development of cancer treatments, lack of standardized and physiologically-relevant *in vitro* testing platforms limit the rapid and early screening of anti-cancer agents. A major barrier in this endeavor, is the complex interplay between the tumor microenvironment and host immune response and lack of predictive biomarkers for clinical benefit. To tackle this challenge, we have developed a dynamic-flow based three-dimensionally (3D) bioprinted vascularized breast tumor model, responding to chemo and immunotherapeutic treatments. Heterotypic tumor spheroids, comprising metastatic breast cancer cells (MDA-MB-231), human umbilical vein endothelial cells (HUVECs) and human dermal fibroblasts (HDFs), precisely bioprinted at pre-defined distances from a perfused vasculature, exhibited tumor angiogenesis and cancer invasion. Proximally bioprinted tumors (∼100 μm) exhibited enhanced capillary sprouting, anastomosis to perfused vasculature and increased cancer cell migration as compared to distally bioprinted spheroids (∼500 μm). Proximally bioprinted tumors treated with varying dosages of doxorubicin for 72 h enabled functional analysis of drug response, wherein, tumors portrayed a dose-dependent drug response behavior with ∼70% decrease in tumor volume for 1 μM dose. Additionally, a cell based immune therapy approach was explored by perfusing HER2-targeting chimeric antigen receptor (CAR) modified CD8+ T cells for 24 or 72 h through the central vasculature. Extensive CAR-T cell recruitment to the endothelium and substantial T cell activation and infiltration in the tumor site, resulted in ∼70% reduction in tumor growth for high CAR treatment densities, after 72 h of treatment. The presented 3D model paves the way for a robust, precisely fabricated and physiologically-relevant 3D tumor microenvironment platform for future translation of anti-cancer therapies to personalized medicine for cancer patients.

**One Sentence Summary:** A physiologically-relevant 3D bioprinted perfusable vascularized tumor model capable of dynamic screening of chemo and immunotherapeutics.

## INTRODUCTION

Cancer remains one of the leading causes of mortality worldwide, accounting for approximately 10 million deaths in 2020 alone *(1)*. Despite remarkable advances in cancer treatment modalities, dearth of physiologically-relevant pre-clinical diagnostic platforms limits the successful clinical translation of anti-cancer therapeutics *(2)*. Genetic and epigenetic heterogeneity of a tumor microenvironment and the underlying immune-cancer interactions are critical aspects in determining effective therapeutic response *(3, 4)*. Even though the field of immunotherapy has witnessed tremendous growth in checkpoint inhibitor-based immunotherapy against PD-1, PD-L1 and CTLA4 in melanoma *(5)*, non-small cell lung cancer *(6)*, and renal cell carcinoma *(7)*; these treatments have shown promising results in only a subset of patients. Additionally, checkpoint inhibitor therapy for breast cancer is only limited to triple negative breast cancer patients expressing PD-L1 protein *(8)*. Similarly, the adoptive transfer of T cells expressing chimeric antigen receptors (CAR) against tumor associated antigens has produced benefits for hematological diseases but they are less effective for solid tumors *(9)*. Developing effective CAR-T cells specific to breast cancer which can successfully navigate the immunosuppressive breast tumor microenvironment and generate potent anti-tumor response is a major challenge *(10)*. Essentially, inadequate understanding of the inhibitory or stimulatory receptor mediated immune cell activation is a deterrent to developing effective immunotherapy. This necessitates the development of relevant *in vitro* tumor models which would help in disseminating immune-cancer interactions and eventually lead to better clinical translations of CAR-T cell therapy.

Traditional two-dimensional (2D) cell cultures are extremely limited in their ability in recapitulating the heterogeneity of native tumor physiology, cancer progression and metastasis, and animal models are expensive, adhere to strict ethical and legal frameworks that limit the scope and speed of the work, and do not closely represent human pathophysiology. Successful clinical translation of drug testing results obtained from these models is only ∼8% *(11)*. In an attempt to develop more representative platforms, complex 3D models employing multicellular spheroids in conjunction with microfluidics have been employed in studying drug response of tumors under flow *(12)* and endothelial regulation of drug transport in ovarian and lung cancer based tumor spheroids *(13)*. However, these studies have primarily been limited to microfluidic-based devices, which entails a fairly lengthy fabrication process with limited control on localization of tumors and no flexibility of altering their positions with respect to primary vasculature without changing the initial master mold design. Distance of the tumor cells from a perfused blood vessel plays a predominant role in tumor growth as it controls diffusion of oxygen and nutrients, which is critical to tumor survival *(14)*. In contrast to traditional microfluidic techniques, 3D bioprinting offers superior control over spatial location of biologics when creating a perfusable cancer model. Even though 3D bioprinting of tumor cells in biomimetic matrices have been used to study mechanisms of the metastatic cascade by controlling the localization of individual cancer cells from the perfused vasculature, none of these studies incorporated the complexity of the tumor microenvironment, such as angiogenesis, stromal and immune cells *(15, 16)*. The molecular and cellular heterogeneity of the tumor microenvironment controls both tumor behavior as well as host immune response to anti-cancer therapeutics *(17)*. Furthermore, 3D bioprinting of cancer models have been limited to bioprinting of individual cancer cells laden in hydrogels. As such, fabrication of a perfusable vascularized tumor microenvironment for the study of cancer progression, angiogenesis, metastasis, and tumor infiltration by human immune cells has not been realized.

Thus, to develop an *in vitro* 3D model representative of the dynamic tumor niche, herein, for the first time, a physiologically-relevant dynamic-flow based 3D vascularized tumor model has been developed employing aspiration-assisted bioprinting enabling accurate positioning of tumors *(18, 19)*. Heterotypic tumor spheroids, comprising of metastatic breast cancer cells (MDA-MB-231), endothelial cells (HUVECs) and fibroblasts (HDFs) were precisely bioprinted in a collagen/fibrin based biomimetic matrix. Precise control over tumor location also revealed higher invasion index of metastatic MDA-MB-231 cells when bioprinted proximal to the perfused vasculature. A dose dependent reduction in tumor volume following doxorubicin uptake by tumor cells was successfully demonstrated in these 3D bioprinted devices. In addition, we found that CAR-T cells engineered to recognize HER2 protein on MDA-MB-231 cells and eventually induce cancer cell apoptosis upon HER2 recognition. Perfusion of CAR-T cells through the central vasculature revealed extensive recruitment of CAR-T cells to the endothelialized vasculature, T cell infiltration to the tumor site as well as reduction in tumor growth. The presented perfusable tumor model can be harnessed to develop novel anti-cancer therapies by enabling high-throughput screening of drugs or immune modulators, which will eventually help advance the field of precision medicine.

## RESULTS

### Device concept, design and fabrication

In this work, a device was designed with a chamber to contain the biologics with perforations on opposite walls, which would serve as connection ports for perfusion. A stainless-steel wire was inserted through the wall perforations, utilizing a pin-molding technique (Fig. 1A1). About ∼ 60 µL of collagen/fibrin based composite hydrogel loaded with 0.5 million/ml HDFs was carefully deposited to half-fill the device (Fig.1A1). Next, employing aspiration-assisted bioprinting, tumor spheroids were precisely bioprinted at pre-defined distances from the stainless-steel wire (Fig. 1A2, Video S1). Specifically, tumor spheroids were bioprinted at a distance of ∼ 100 µm (proximal) and ∼ 500 µm (distal) from the wire. After bioprinting, the device was then completely filled with additional (∼ 60 µL) deposition of the same hydrogel containing HDFs (Fig. 1A3). After complete crosslinking of the hydrogel (at 37 °C for 30 min), the stainless-steel wire was then gently removed from the device (Fig. 1A4), leaving an open channel. Next, HUVECs were introduced into the channel at a concentration of 25-30 million cells/ml (Fig. 1A5). The following day, the device was connected to an external pump to initiate media flow through the endothelialized channel (Fig. 1A6). The bioprinted tumor spheroids gradually developed angiogenic sprouts under perfusion (Fig. 1A7). Devices were further evaluated for their efficacy in chemo and immunotherapy, as presented later in the paper.

**Figure 1:**
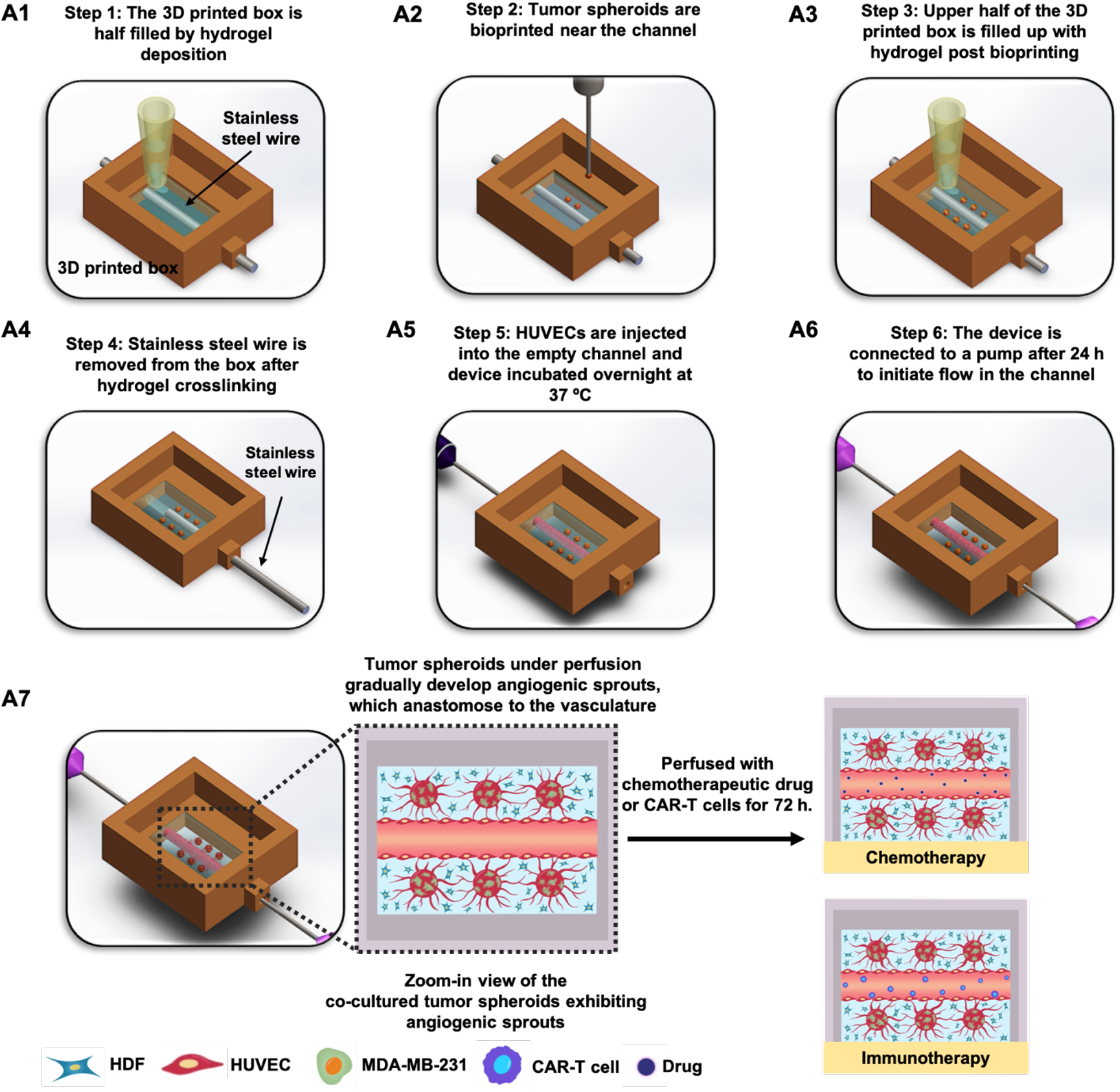
A schematic representation of a 3D perfusable tumor model fabricated using aspiration-assisted bioprinting. (A1-A6) Step 1-Step 6 enumerates the design and work flow followed for successful fabrication of the perfusable device. (A7) Schematic representation of the tumor angiogenesis observed after device is connected to a dynamic flow-based culture. After induction of tumor angiogenesis, the device was utilized for chemo and immunotherapy.

### Evaluation of tumor spheroid and composite hydrogel properties for precise positioning of these spheroids

Aspiration-assisted bioprinting was utilized for precise positioning of tumor spheroids, which involved aspirating tumor spheroids and placing them at desired locations into a biomimetic matrix. Thus, it was crucial to assess the structural and mechanical properties of spheroids for successful bioprinting. In this study, first, we sought to compare three different types of tumor spheroids, homocellular spheroids formed by MDA-MB-231 cells (MDA-MB-231-only), co-cultured spheroids formed by HUVECs and MDA-MB-231 cells (H231), and a third type formed by a combination of HUVECs, MDA-MB-231 cells and HDFs (H231F). These three different types of spheroids exhibited not only distinct morphological differences but also contrasting mechanical properties. MDA-MB-231-only spheroids were unable to withstand the back pressure and experienced a major deformation in their structure due to aspiration (Fig. 2A1). The portion of the spheroid inside the pipette, denoted as the aspiration length, increased with aspiration time for MDA-MB-231-only spheroids (Fig. S1A). Eventually, MDA-MB-231-only spheroids disaggregated and were completely aspirated, indicating their extremely weak mechanical properties and low structural integrity; thus, unsuitable for 3D bioprinting. Next, on repeating the same process for H231 and H231F spheroids, the latter exhibited the highest structural integrity as little to no deformation occurred in H231F spheroids even after 10 min of exposure to back pressure. On measuring the elastic modulus of all spheroid types, H231F exhibited the highest average modulus, ∼160 Pa, as compared to H231, which had an average modulus of ∼105 Pa (Fig. 2A2). We were unable to get a definite elastic modulus value for MDA-MB-231-only spheroids as they completely lost their structural integrity during elastic modulus measurement. Ultrastructural images of all spheroid types revealed a dense and compact morphology for H231F as compared to H231 and MDA-MB-231-only spheroids (Fig. 2A3). Thus, owing to their superior mechanical and structural integrity, and heterotypic nature mimicking tumor microenvironment, H231F spheroids were preferred for 3D bioprinting experiments. Immunostaining confirmed the presence of all cell types utilized, where, HDFs, identified by PDGFRα staining, were located at the peripheral region of the spheroid body, GFP^+^ MDA-MB-231 cells were dispersed throughout the inner and outer regions of the spheroid body, while tdTomato^+^ HUVEC occupied a majority (Fig. 2B).

**Figure 2:**
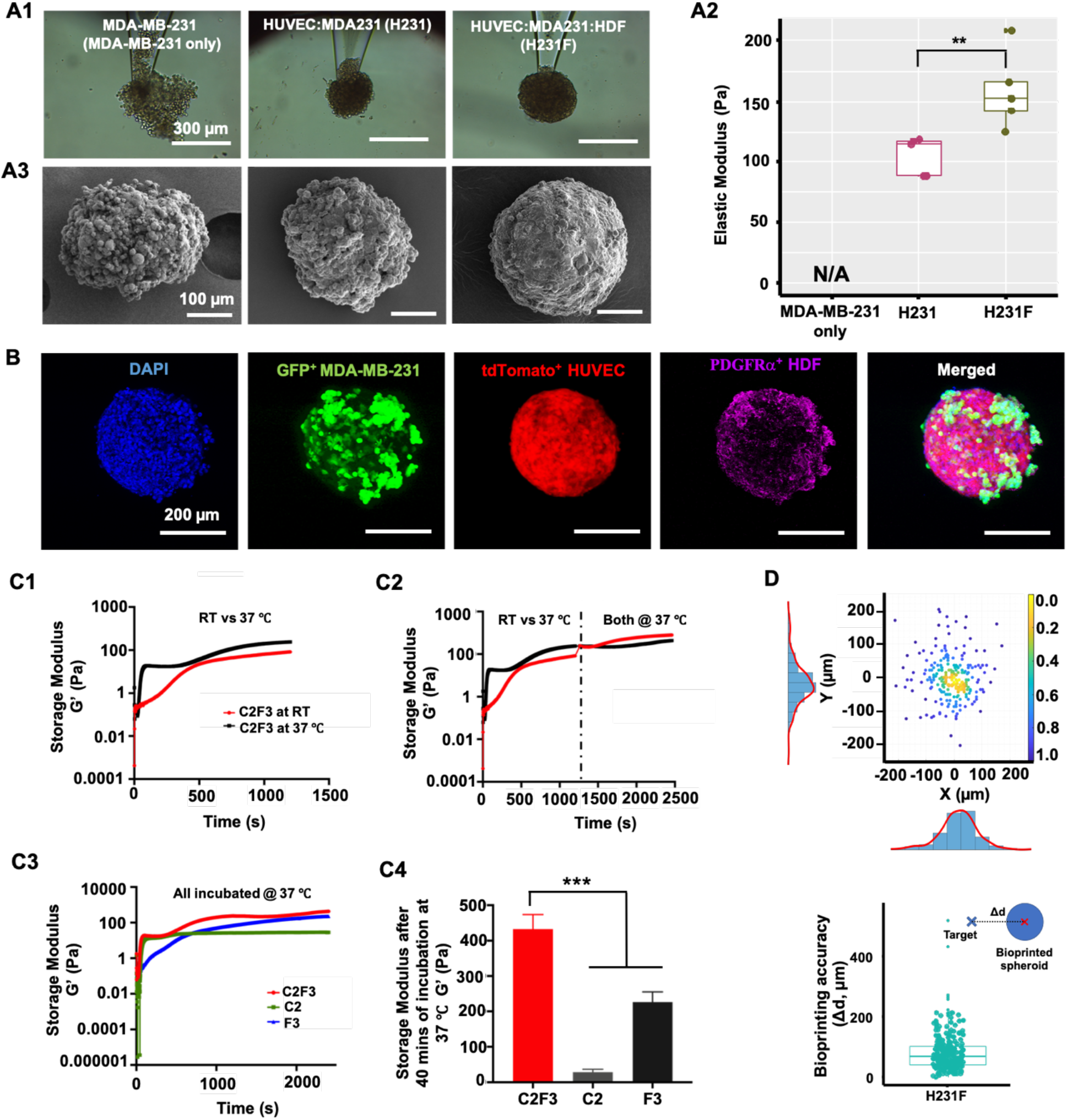
Evaluation of spheroid and hydrogel properties for aspiration-assisted bioprinting. Three different tumor spheroids were used to compare the mechanical and structural properties. Tumor spheroids used were spheroids formed by only MDA-MB-231 cells (MDA-MB-231-only), combination of HUVECs and MDA-MB-231s (H231) and combination of HUVECs, MDA-MB-231 and HDFs (H231F). (A1) Tumor spheroids were aspirated with a glass micropipette for 10 min to analyze their structural integrity under aspiration. (A2) Graphical representation of elastic modulus measured for the tumor spheroids (*n*=3 for all, p^***^ < 0.001, p^**^< 0.01, p^*^< 0.5). (A3) SEM images of tumor spheroids exhibiting distinctly different surface topographies. (B) Fluorescent images of the composite H231F tumor spheroids displaying all three different cell types, namely GFP^+^ MDA-MB-231 cells, tdTomato^+^ HUVECs and PDGFRα^+^ HDFs. DAPI (blue) stained all cell nuclei. (C1) Graphical representation of change in storage modulus over time of the composite hydrogel (C2F3) incubated at room temperature (RT) and 37 °C for 20 min. (C2) Graphical representation of change in storage modulus over time for C2F3 incubated at RT and 37 °C for 20 min and then incubated at only 37 °C. (C3) Graphical representation of change in storage modulus over time for C2F3 vs. its precursor, C2 and F3, all incubated at 37 °C for 40 min. (C4) Graphical representation of storage modulus for C2F3 as compared to its precursors, C2 and F3, all incubated at 37 °C for 40 min. (D) Graphical representation of bioprinting positional accuracy for spheroids bioprinted at defined X, Y locations (graph on top) and a measured drift calculated between the target and actual location (bottom graph) (*n*=250).

Collagen and fibrin, both being abundantly present at a native tumor site, were incorporated in our study as the bioprinting substrate. Building on our previous study *(20)*, we incorporated 2 mg/ml collagen along with 3 mg/ml fibrinogen in this study (C2F3), to build the tumor microenvironment. In order to precisely position H231F spheroids in the composite C2F3 hydrogel, it was essential to realize the entire bioprinting process prior to complete crosslinking of C2F3. Bioprinting inside a crosslinked hydrogel would result in irreversible gel deformation as C2F3 did not possess self-healing properties. Thus, to understand the rheological changes in C2F3 during bioprinting performed at room temperature (RT, 22 °C), a time sweep test was performed, first at room temperature, and then at 37 °C. A gradual increase in storage modulus was observed for C2F3 incubated at RT, for the first ∼ 300 s (5 min) (Fig. 2C1). In contrast, a sharp increase in storage modulus was observed for C2F3 incubated at 37 °C, in the first few seconds itself. The storage modulus eventually acquired a constant value of ∼100 Pa in ∼ 20 min at both RT and 37 °C. This implied that incubating C2F3 at RT offered a time window of ∼ 5 min to complete the bioprinting process before onset of crosslinking. Further, to understand if the rheological properties of C2F3 altered when finally incubated at 37 ^°^C after bioprinting, we performed a similar time sweep test at 37 ^°^C after the initial 20 min run at RT (Fig. 2C2). The storage modulus for C2F3 eventually recovered and reached a mean value of ∼ 400 Pa, which closely matched that of C2F3 incubated at 37 ^°^C throughout. Next, the trend in the storage modulus of the precursors C2 and F3 were compared to that of C2F3 by conducting a time sweep test at 37 ^°^C (Fig. 2C3). C2F3 exhibited a significantly high storage modulus, ∼ 400 Pa, as compared to the precursors C2, ∼50 Pa, and F3, ∼200 Pa (Fig. 2C4).

On optimizing both the matrix and tumor spheroid properties required for 3D bioprinting, bioprinting accuracy was also determined for H231F spheroids, which was measured to be ∼50 µm (Fig. 2D). Additionally, the circularity of spheroids before and after bioprinting was also determined to be similar (∼ 0.5) indicating that bioprinting had minimal effect on spheroid morphology (Fig. S1C).

### 3D bioprinting of tumor spheroids at varying distances from the perfusable vasculature

Employing aspiration-assisted bioprinting, H231F spheroids, were bioprinted at pre-defined distances from the perfusable vasculature to analyze the effect of distance on tumor angiogenesis and cancer invasion. To confirm the barrier function of the engineered vasculature, the diffusional permeability was evaluated and compared with respect to non-endothelialized channels (empty), Non-endothelialized channels exhibited a significantly higher diffusion of fluorescein isothiocyanate (FITC)-conjugated dextran as compared to the endothelialized channels (Fig. 3A1). Diffusional permeability for empty channels in C2F3 had a mean value of ∼ 7×10^−5^ cm/s as compared to ∼ 2×10^−5^ cm/s, for endothelialized channels. Additionally, CD31 staining of HUVECs in the endothelialized channel exhibited their uniform lining throughout the vasculature (Fig. 3A2). H231F spheroids were bioprinted either proximal (at a distance of ∼ 100 µm) or distal (∼ 500 µm) to the endothelialized channel (vasculature). To understand the effect of distance from the vasculature on both tumor angiogenesis and metastasis, three different types of devices were bioprinted. In the first set of devices, all spheroids were bioprinted at an average distance of ∼ 100 µm on both sides of the vasculature (proximal, Fig. 3B1). Next, spheroids were bioprinted at an average distance of ∼ 100 µm on one side (proximal) and ∼ 500 µm (distal) on the other side of the vasculature, as shown in Fig. 3B2. The third device included all spheroids bioprinted at ∼ 500 µm distance (distal, Fig. 3B3). Tumor angiogenesis was induced in all these devices under perfusion, maintaining a perfusion speed of 0.7 μL/min. Optimal sprout formation and maintenance of endothelial barrier was observed under this flow rate. The angiogenic sprouts developed hollow capillaries (Fig. S2A) and vascularized the tumor spheroids (Figs. 3C1-C3, Movie S2). For the proximal spheroids, sprouts further anastomosed and connected to the main vasculature, representative of *in vivo* physiology. It was interesting to observe that the extent of tumor angiogenesis with regards to the total vessel length and vessel branching varied with distance of spheroids from the perfused vasculature. A higher sprouting density was observed for proximal tumors as compared to the distal spheroids (Figs. 3C1-C3). The total vessel length for the proximal spheroids had an average value of ∼2.5 mm as compared to ∼1.2 mm for the distal spheroids (Fig. 3D1). Additionally, the total number of junctions was approximately 2-fold higher for the proximal spheroid as compared to the distal ones (Fig. 3D2). Additionally, the sprouts formed in the distal spheroids were unable to connect to the perfused vasculature. The distance of spheroids from the vasculature also affected cancer invasion. For proximal tumor spheroids, cancer cells proliferated and were observed to invade into the vasculature (Fig. S2B, Movie S3). Spatial location of tumor spheroids also affected cancer invasion. MDA-MB-231 cells invaded into its surrounding matrix for both the proximal and distal spheroids. However, the area occupied by proximal spheroids was ∼ 1.5-fold higher than the area occupied by distal spheroids (Fig. 3D3).

**Figure 3:**
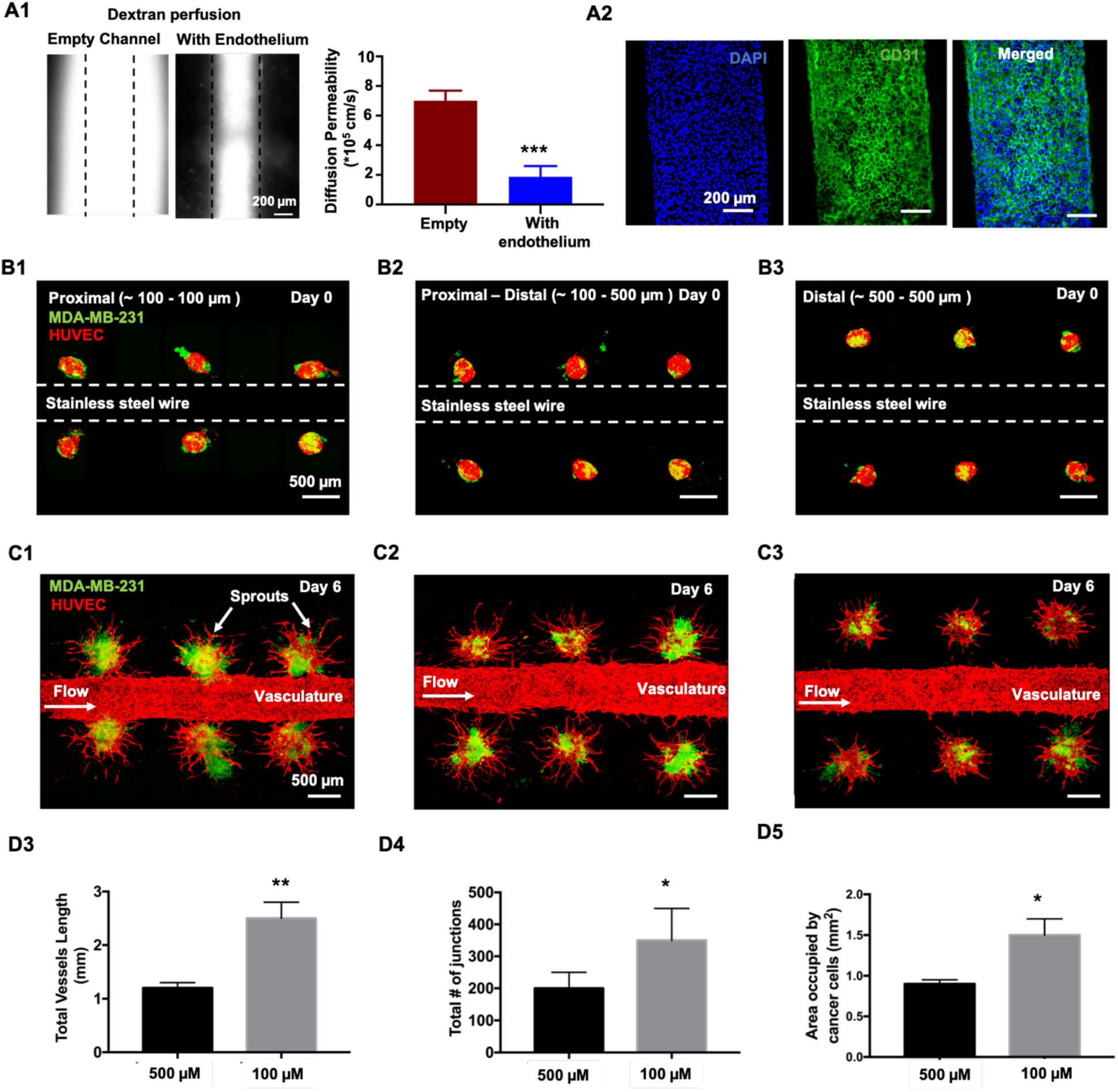
(A1) Fluorescent images of dextran perfusion in empty and endothelialized channels and their corresponding permeability values (A2) Confocal images of endothelialized channels stained for CD31 and DAPI (4′,6-diamidino-2-phenylindole). (B1-B3) Devices with tumor spheroids bioprinted at a proximal only (distance of ∼ 100 µm), proximal-distal combined (distance of ∼ 100 µm and ∼ 500 µm) and distal (∼500 µm) from the stainless-steel wire. (C1-C3) Fluorescent images of bioprinted devices after six days of perfusion exhibiting tumor angiogenesis. (D3–D5). Graphical representation of total vessels length, total number of junctions and area occupied by cancer cells as a function of distance from the central vasculature (*n*=3 for all, p^***^ < 0.001, p^**^ < 0.01, p^*^ < 0.05).

### Effects of chemotherapeutic drug doxorubicin, on tumor growth under a dynamic flow-based culture

In this study, doxorubicin, an anthracycline based chemotherapeutic drug *(21)*, commonly used for treating breast cancer was tested to understand drug response in a dynamic flow-based microenvironment. We first tested doxorubicin on tumor spheroids under two different static culture conditions prior to perfusing it through devices. These static cultures included (i) free-standing spheroids directly exposed to doxorubicin and (ii) C2F3 encapsulated spheroids exposed to doxorubicin. Doxorubicin concentration was varied from 0.0001 to 100 µM and spheroids were treated with it for a period of 72 h. A non-treated group was used as control, containing only vehicle (0.1% DMSO). Herein, we tested the effect of this drug on two different types of tumors, including MDA-MB-231-only and H231F spheroids. In particular, we aimed to assess if the presence of other cell types affected the drug response. As depicted in fluorescent images in Fig. S3A1, MDA-MB-231-only spheroids showed a significant reduction in GFP intensity for doxorubicin concentrations ranging from 1 to 100 µM. There was also a reduction in spheroid size for this concentration range, evident from the fluorescent images. Viability of spheroids was measured after a 72 h doxorubicin treatment to understand their dose-response behavior (Fig. S3A2). An IC_50_ value of ∼ 0.51 µM was obtained for MDA-MB-231-only spheroids. Within a 95% confidence interval (C.I.), the IC_50_ range was predicted to be _∼_ 0.3 – 0.85 µM for MDA-MB-231-only spheroids. Similarly, co-cultured H231F spheroids were also treated with varying concentrations of doxorubicin for 72 h. As shown in Fig. S3B1, there was a reduction in GFP intensity for 1 - 100 µM. An IC_50_ value of ∼ 0.65 µM was obtained from the dose response curve (Fig. S3B2). Additionally, within a 95% C.I., the IC_50_ range for H231F was predicted to be ∼ 0.4 – 1.1 µM.

To understand the effect of extracellular matrix (ECM) on drug response of tumors, we encapsulated MDA-MB-231-only and H231F spheroids in C2F3, without any HDFs in the matrix. These spheroids were first cultured for a period of 3 days to allow them to grow in C2F3 and then varying concentrations of doxorubicin was introduced to these cultures for 72 h. As shown in Fig. S4, there was a significant reduction in GFP intensity for 10 and 100 µM doxorubicin treated spheroids, indicating extensive cell death. For 0.001 to 1 µM, there was also a reduction in tumor size and invasion, as compared to the non-treated or vehicle control groups. Additionally, we obtained an IC_50_ value of ∼ 0.22 µM with an IC_50_ range of 0.12-0.3 µM (95% C.I.), which was much lower than the MDA-MB-231-only group in previous experiments, where free-standing tumors were directly exposed to doxorubicin. This observation suggested that tumor-ECM interactions could affect drug outcomes. Interestingly, when H231F spheroids were encapsulated in C2F3 and treated with doxorubicin for 72 h, tumor viability ranged from ∼88-97% for 0.0001 - 0.1 µM (Figs. S4B1-B2). Fluorescent images in Fig. S4B1 revealed that GFP intensity was most affected by 1, 10, and 100 µM concentrations. We also obtained a higher IC_50_ value of ∼ 0.9 µM as compared to 0.2 µM for the MDA-MB-231-only spheroids encapsulated in H231F. Similarly, an IC_50_ range of ∼ 0.6 - 1.3 µM (95% C.I.) was obtained for H231F spheroids encapsulated in C2F3.

After analyzing the effect of doxorubicin on static cultures, we introduced it at varying concentrations in perfusable devices. For this study, all spheroids were bioprinted at a distance of ∼ 100 µm (proximal) from the perfused vasculature. Spheroids cultured under flow for 3 days were observed to induce angiogenic sprouting which further anastamosed with the central vasculature. Doxorubicin (0 - 100 µM), diluted in EGM-2MV media, was perfused through the central vasculature. We were unable to use the Alamar Blue viability assay on these devices, as the presence of HDFs in C2F3 dominated the assay output and it was not sensitive enough to capture the changes in MDA-MB-231 viability. Thus, we used Caspase assay to detect early onset of apoptosis, after 24 h of doxorubicin treatment and imaging techniques to capture the tumor fate after 72 h of doxorubicin treatment. To analyze doxorubicin uptake by cancer cells under perfusion and co-localization of the drug in cell nuclei, doxorubicin was excited at 780 nm. The tumor core was imaged to understand if the drug had diffused to MDA-MB-231 cells at that location. There was an extensive uptake and localization of doxorubicin in the tumor core for both 10 and 100 µM treatments, identified by the doxorubicin auto-fluorescence signals (Fig. 4A1). Widespread cellular debris in tumor core and significantly less density of GFP^+^ MDA-MB-231 cells were observed for these two above-mentioned concentrations. For the 1 µM treatment, doxorubicin auto-fluorescence was detected in tumor core but much lower as compared to 100 and 10 µM treatments. Post 1 µM treatment, doxorubicin was not detected in the tumor core even while using a higher laser power, for rest of the drug concentrations. As doxorubicin induces cellular apoptosis, we sought to understand Caspase 3/7 activation after 24 h of doxorubicin perfusion in devices. As shown in Fig. 4A2, Caspase 3/7 activity was observed to be ∼ 5-7 folds higher for 10 and 100 µM treatments, as compared to the non-treated group. Additionally, for the 1 µM treatment, Caspase 3/7 activity was 3-fold higher than that for the non-treated group. For 0.1 to 0.0001 µM treatments, no significant difference in Caspase activity was observed as compared to the non-treated group.

**Figure 4:**
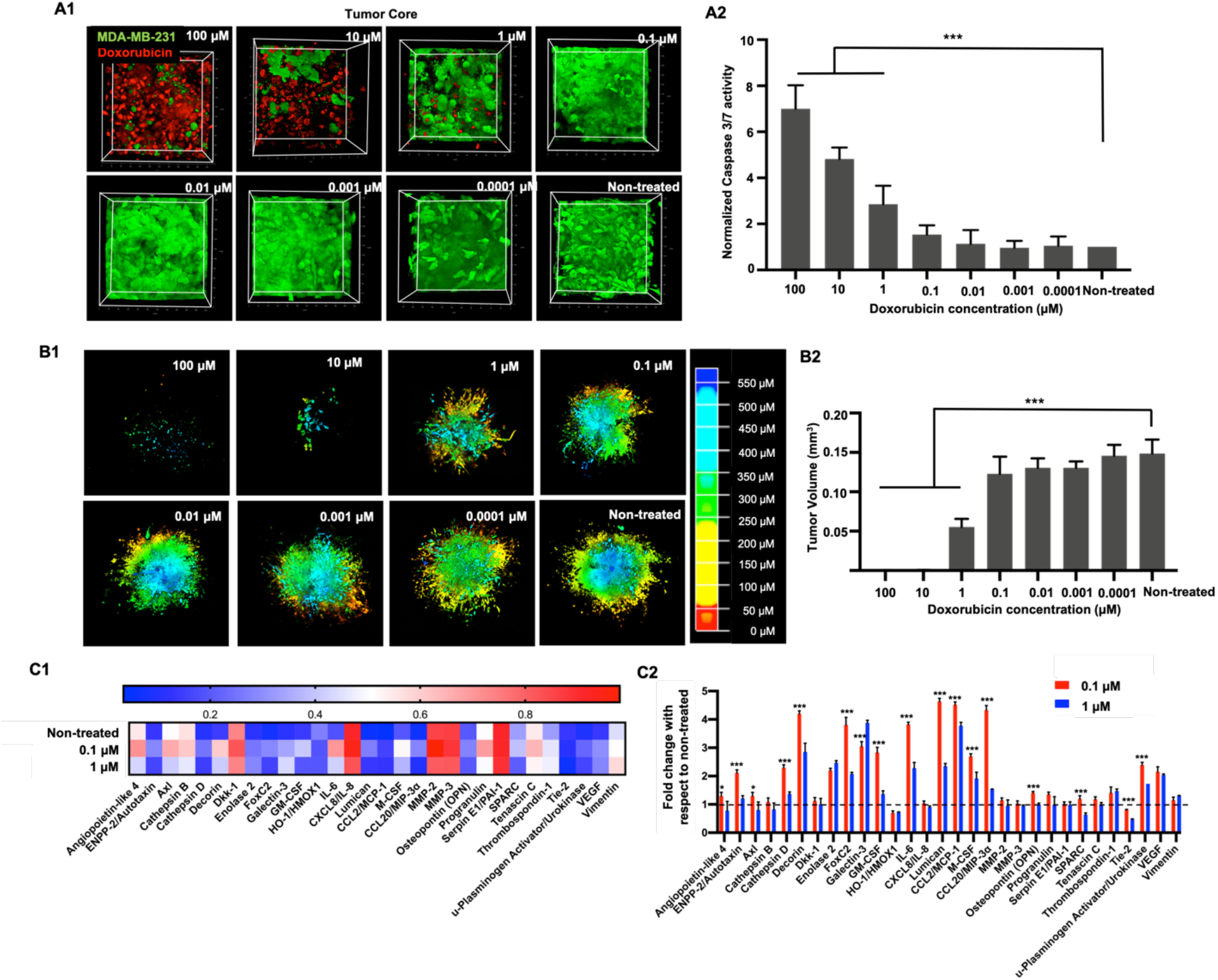
(A1) 3D Reconstruction of the tumor core showing GFP^+^ MDA-MB-231 cells and auto-fluorescent doxorubicin co-localized in cell nuclei. Doxorubicin concentration was varied from 100 to 0.0001 µM. (A2) Graphical representation of normalized caspase 3/7 activity after 24 h of the doxorubicin treatment. (B1) 3D Reconstruction of the entire tumor remaining after 72 h of doxorubicin treatment. The tumors were depth coded to represent the entire tumor volume. (B2) Graphical representation of tumor volumes after 72 h of doxorubicin perfusion. (C1) Heatmap of selected proteins from the human oncology array performed on perfusates collected from devices treated with 1, 0.1 and 0 µM (non-treated) Doxorubicin. (C2) Graphical representation of fold change in protein expression for treated devices as compared to non-treated devices. The dotted line at 1 through the bar graph helps identify protein expressions, which had values close or equal to the non-treated group. Thus, values higher than 1 represented an increase in the protein expression and values lower than 1 depicted decrease in expression as compared to the non-treated group (*n*=3 for all, p^***^< 0.001, p^**^ < 0.01, p^*^ < 0.05).

After perfusing doxorubicin for 72 h, GFP^+^ MDA-MB-231 cells remaining in the entire tumor volume was imaged for all concentrations. As shown in Fig. 4B1, 10 and 100 µM treatments were extremely cytotoxic, resulting in only a few MDA-MB-231 cells remaining viable post treatment. This was also observed from our previous static cultures. 1 µM treatment was also cytotoxic to MDA-MB-231 cells, resulting in fewer GFP^+^ cells remaining. Doxorubicin cytotoxicity was observed to be waning from 0.000 to 0.1 µM. As compared to the non-treated group, where the tumor spanned a depth of ∼ 600 µm, a significantly smaller tumor was found for 10 and 100 µM treatments. The 1 µM treatment was also observed to be less dense as compared to other concentrations. On quantifying the tumor volumes, as expected, a significantly low volume (0.0004 mm^3^) was found for 10 and 100 µM treatments. For the 1 µM treatment, a mean tumor volume of ∼ 0.05 mm^3^ was obtained. For 0.0001– 0.1 µM, tumor volumes were determined to range from ∼ 0.12 to 0.15 mm^3^, which were close to the non-treated group (∼ 0.15 mm^3^).

In order to understand the effect of doxorubicin on common cancer-related protein expressions, a proteome array containing 84 human oncoproteins was used to detect protein expressions in treated devices. The experimental groups chosen for this analysis included 0.1 µM, 1 µM, and non-treated (control) perfusion devices. As 10 and 100 µM treatments resulted in extensive cell death in static as well as perfusion cultures, 1 µM was chosen as the next ‘high’ concentration to be analyzed. Additionally, as the 0.1 µM treatment resulted in comparable tumor volumes after 72 h of treatment with respect to the non-treated group, 0.1 µM would represent a ‘low’ concentration. Fig. S5 portrays a heatmap of all 84 oncoproteins from which a group of 29 proteins were identified having significantly high and variable expression among the three experimental groups. As shown in the heatmap in Fig. 4C1, Angiopoietin-like 4, Axl, Capthepsin-B, Dkk-1, IL-8, MMP-2, MMP-3, Progranulin, Serpin E1, Tenascin C and Vimentin were found to be comparatively higher in expression (> 0.5) as compared to other proteins, for all treated and non-treated groups. Overall, on comparing the two doxorubicin treated groups, the 0.1 µM treatment exhibited expressions values higher than the non-treated group for some proteins. Thus, to further understand how the protein expression in the treated groups varied from the non-treated ones, fold change of protein expression was calculated with respect to the non-treated group (Fig. 4C2). We found comparable expressions of Angiopoietin-like 4, Axl, Capthepsin-B, Dkk-1, CXCL8/IL-8, MMP-2, MMP-3, Progranulin, Serpin E1, Tenascin C and Vimentin, for all the three groups, irrespective of doxorubicin treatment. There was a 2-4-fold increase in expression levels of Decorin, Enolase-2, FoxC2, Galectin-3, IL-6, Lumican, CCL2/ MCP-1, M-CSF, GM-CSF Osteopontin (OPN), uPA, and VEGF for both doxorubicin treated groups (0.1 and 1 µM), as compared to the non-treated group. Additionally, the 0.1 µM treatment exhibited increased expression of ENPP-2/ Autotaxin, Cathepsin D, and CCL20, while all of these were similar in expression for the 1 µM treatment, when compared to the non-treated group. In contrast, a 0.5-0.7-fold decrease in the expression of HMOX1 was observed for both 0.1 and 1 µM treated groups, and a decrease in SPARC and Tie-2 expression was observed for the 1 µM treatment group.

### Tumor growth during dynamic flow-based culture with anti HER2 CAR-T cells

We next asked whether this 3D solid tumor model could be purposed to study or develop novel cell-based immune therapy strategies *in vitro*. Accordingly, breast tumors were treated with chimeric antigen receptor expressing CD8+ human T cells engineered to express anti-human epidermal growth factor receptor 2 (HER2) in conjunction with chimeric antigen receptor (CAR) synthetic genes as described *(22), (23), (24)* and as in methods. Anti HER2 targeting CAR-T cells, tested under *in vitro* static cultures on homotypic MDA-MB-231 and heterotypic H231F tumors, exhibited a dose dependent decrease in GFP intensity of MDA-MB-231 cells, over 3 days of culture (Figs. S6A1, S7A1). Both tumor types exhibited ∼60-100% reduction in GFP intensity with increasing anti HER2 CAR-T cell treatment density (Figs. S6A3, S7A3). Whereas heterotypic H231F tumors exposed to anti CD19 CAR T cells did not exhibit any significant decrease in GFP intensity (Fig. S7A2). In contrast, anti CD19 CAR-T cell treatment of homotypic MDA-MB-231 tumors resulted in ∼60-80% decrease in GFP intensity for treatment densities higher than 2xT (Figs. S6A2, S7A3). Following *in vitro* static cultures, CAR-T cells were perfused through the engineered vasculature to analyze the cancer-immune interaction in a dynamic-flow based culture (Movie S4). After 24 h, CAR-T cells were found adhering to the central vasculature, through which they were perfused (Fig. 5A1). The density of CAR-T cells adhering to the endothelium increased after 72 h of perfusion, which indicated an inflammatory microenvironment (Fig. 5A2). Additionally, CAR-T cells were also found to have infiltrated to the tumor site, probably via the anastomosed capillaries and the porous C2F3 matrix and CAR-T cells were found located near the sprouts as well as inside the tumor associated capillaries (Fig. 5A3).

**Figure 5:**
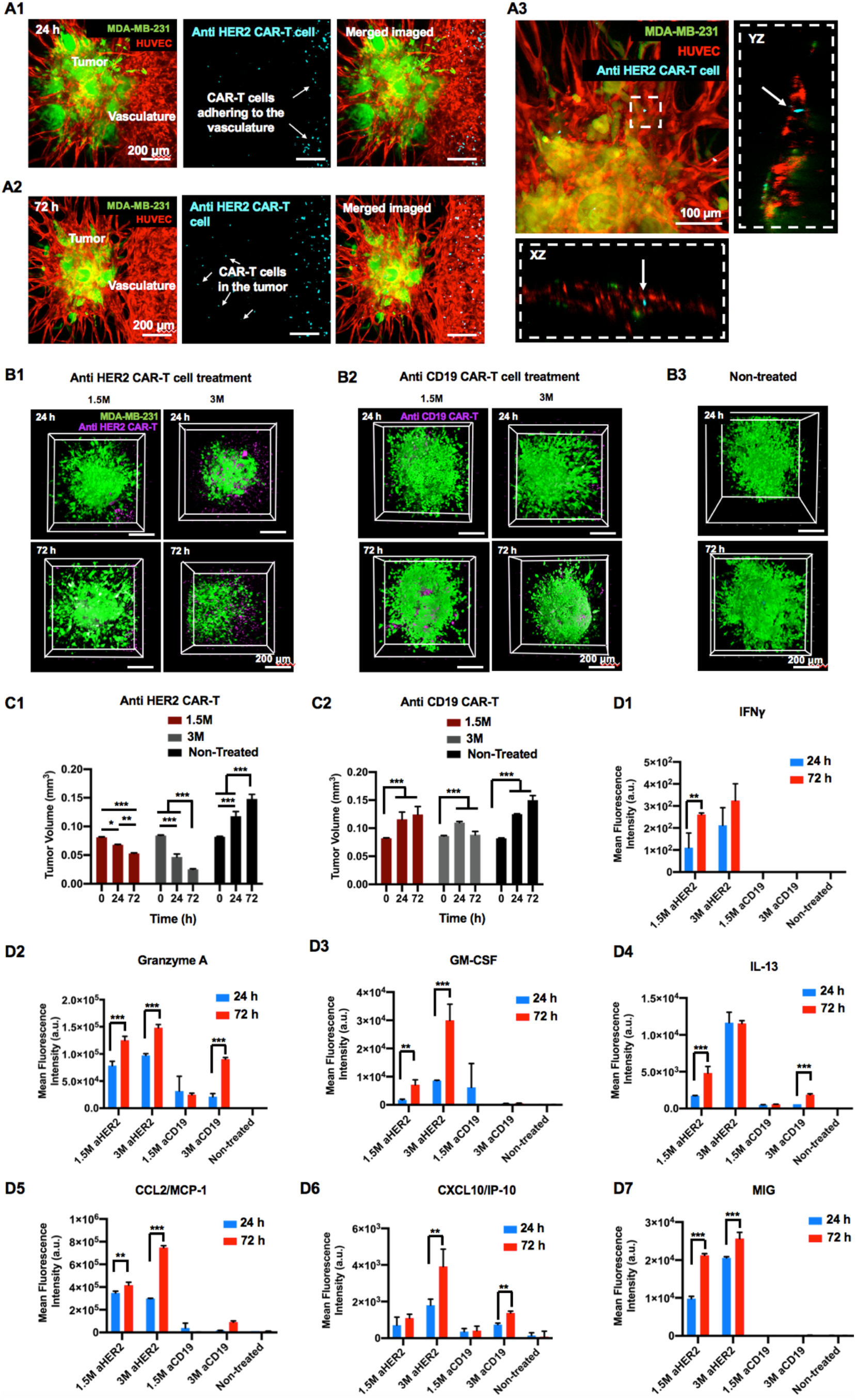
Culture of engineered CAR-T cells in a 3D bioprinted perfusable and vascularized tumor microenvironment. (A1) Fluorescent images of devices perfused with anti HER2 CAR-T cells for 24 h. CAR-T cells were labelled with cell tracker violet (CTV) and observed to adhere to HUVECs in the perfused vasculature. (A2) Fluorescent images of devices perfused with anti-HER2 CAR-T cells for 72 h. CAR-T cells were observed to have infiltrated to the tumor site. (A3) Orthogonal projection of anti HER2 CAR-T cells inside a capillary (denoted by white arrows). 3D Reconstruction of tumor cores illustrating GFP^+^ MDA-MB-231 cells and CTV-labelled (B1) anti-HER2 CAR-T cells and (B2) anti-CD19 CAR-T cells. CAR-T cell treatment density was varied as 1.5 and 3 million active CAR-T cells. Perfusion was carried out for 24 h or 72 h for both concentrations. (B3) 3D Reconstruction of tumor cores illustrating the GFP^+^ MDA-MB-231 cells for non-treated (control) groups. (C1) Graphical representation of tumor volumes after 24 or 72 h of (C1) anti-HER2 CAR-T cell and (C2) anti CD19 CAR-T cell perfusion (*n*=3 for all, p ^***^< 0.001, p^**^ < 0.01, p^*^ < 0.05). Graphical representation of the mean fluorescence intensity of the cytokines and chemokines secreted after 72 h of CAR-T cell perfusion including (D1) IFNγ, (D2) Granzyme A, (D3) GM-CSF, (D4) IL-13, (D5) CCL2/ MCP-1, (D6) CXCL10/ IP-10 and (D7) MIG (*n*=3, p^*^<0.05, p^**^<0.01, p^***^<0.001).

To better understand the effect of CAR-T cell treatment on the bioprinted tumor microenvironment, two treatment densities, comprising of 1.5 million (1.5M) and 3 million (3M) CAR-T cells were perfused for a period of 24 or 72 h. For both treatment densities, CAR-T cells had infiltrated to the tumor site, with a higher density of T cells in the tumor core observed for the 3M group (Figs. 5B1, B2). For the 3M treatment density, presence of CAR-T cells in the tumor core was visibly higher after 72 h of perfusion for the anti HER2 treatment as compared to the anti CD19 treatment. Under this dynamic flow-based culture, anti HER2 CAR-T cells suppressed the tumor growth as compared to the non-treated group (Figs. 5B1-B3). On comparing the tumor volume (∼0.081 mm^3^) prior to onset of treatment (0 h), tumors under 1.5M anti HER2 CAR-T cell treatment for 24 h had a ∼16% reduction in volume (∼0.068 mm^3^) and a further ∼22% reduction (0.053 mm^3^) after 72 h treatment (Figs. 5C1, S8A). Increasing the treatment density to 3M resulted in a significant ∼44% (∼0.047 mm^3^) and a further ∼46% (∼0.025 mm^3^) decrease in tumor volume after 24 and 72 h CAR-T cell perfusion, respectively (Figs. 5C1, S8A). In contrast, as compared to the tumor volume before the treatment (0.082 mm^3^), perfusing 1.5M anti-CD19 CAR-T cells resulted in a ∼41% (0.116 mm^3^) increase in tumor growth after 24 h and a further increase of ∼7.5% (0.124 mm^3^) in the next two days (72 h) (Figs. 5C2, S8B). Increasing the anti-CD19 CAR-T cell treatment density to 3M resulted in a ∼28% (0.11 mm^3^) increase in tumor volume after 24 h of CAR-T cell perfusion and a subsequent ∼20% decrease (0.088 mm^3^) in tumor growth after 72 h of perfusion (Figs. 5C2, S8B). Tumor volumes of non-treated tumors increased by ∼80% (∼0.15 mm^3^) in volume after 72 h of perfusion (Figs. 5C1-C2, S8C).

Cytokines and chemokines, secreted as an aftermath of the cancer-immune interactions, were assessed from the culture supernatants collected from the last 24 h of the 72-h-culture with CAR-T cells. Increased expression of interferon-gamma (IFNγ) was only observed after anti HER2 CAR-T cell perfusion, for both 1.5M and 3M treatments, which was indicative of HER2 specific CAR-T cell activation (Fig. 5D1). Subsequent cytotoxic Granzyme A secretion was also observed to be higher for the anti HER2 CAR-T cell treatment, as compared to anti CD19 treatment (Fig. 5D2). Additionally, cytokines GM-CSF and IL-13, and chemokines CCL2 MCP1, IP-10, and MIG were all detected in higher quantities in the culture perfusates of anti HER2 CAR-T cell treated tumors as compared to anti CD19 CAR-T cell treated counterparts (Figs. 5D3-C7). Furthermore, expression of all cytokines and chemokines were higher after 72 h of CAR-T cell perfusion as compared to 24 h, which indicated enhanced immune activation and immune response over time. Enhanced secretion of these immune activators was also observed under *in vitro* static culture conditions for both homotypic MDA-MB-231 and heterotypic H231F tumors (Figs. S6, S7). Specifically, expression of IFNγ, Granzyme A, GM-CSF, and IL-13 were higher with anti HER2 CAR-T cell treatment as compared to anti CD19 CAR T control condition (Figs. S6B1-B4, S7B1-B4). In contrast, chemokines CCL2 MCP1, IP-10 and MIG were equally expressed in *in vitro* static cultures after both anti HER2 and anti CD19 CAR T-treatments, which indicated higher degree of non-specific cancer immune interactions in static cultures (Figs. S6B5-B7, S7B5-B7).

## Discussion

This study demonstrates, for the first time, a 3D bioprinted vascularized tumor model responding to both *in vitro* chemotherapy as well as immunotherapeutic treatments. Employing aspiration-assisted bioprinting, heterotypic tumors comprising of endothelial cells, cancer cells and fibroblasts were precisely bioprinted at desired locations from a main vasculature in a composite collagen/fibrin biomimetic matrix. Constant flow of media supplemented with growth factors, at low shear stress through the main vasculature, eventually induced tumor angiogenesis and aided cancer metastasis in tumors bioprinted proximal or distal to the vasculature. Low shear stress supported both angiogenesis as well as maintained endothelium integrity over time, as high shear stresses have shown to be inhibitory to sprouting *(25)*. The incorporation of fibroblasts in spheroids not only rendered them mechanically and structurally suitable for bioprinting, it also helped to recapitulate the heterotypic nature of native tumor microenvironment. The tumor microenvironment is often considered to possess a ‘reactive stroma’ which is typically associated with the presence of several cellular (fibroblasts, endothelial and immune cells) and acellular ECM components (enhanced Type I collagen and fibrin deposition) *(26)*, all of which distinctly differ from their quiescent counterparts in homoeostatic tissues and plays an astounding role in cancer progression *(27)*.

Employing 3D bioprinting to precisely position tumors in a tumor stroma at pre-defined distances could potentially enable us to study the role of paracrine signaling in tumorigenesis. In this study, spheroids bioprinted proximal or distal to the main vasculature and subjected to dynamic flow-based conditions, exhibited differential tumor angiogenesis based on the location of the tumor. Tumors bioprinted proximal to the central vasculature developed longer and denser vessels with a higher junction density. Enhanced diffusion of nutrients through the main vasculature towards the proximal tumors and subsequent local accumulation of growth factors and cytokines, such as VEGF-A, basic fibroblast growth factor, and platelet-derived growth factor-BB, could enhance vessel formation for tumors located close to the central vasculature *(28)*. This could also be attributed to the probable upregulation of local matrix metalloproteinases (MMPs) near the perfused vasculature, which contributes to greater matrix degradation, thus enabling HUVECs in the proximal tumors to develop longer vessels *(29)*. Moreover, these results also suggest that HUVECs associated with the tumor are capable of sensing metabolic and physiological changes in the surrounding stroma, which eventually activates the angiogenic signaling cascade *(30)*. Interestingly, the distance of spheroids from the perfused vasculature also affected cancer invasion.

Distal tumors exhibited decreased invasion as compared to proximal tumors and this phenomenon corroborates the fact that cancer cells can alter their behavior based on extracellular stimuli originating from the surrounding microenvironment *(31)*. Being close to the perfused vasculature, there is probably increased expression of urokinase-plasminogen activator (uPA) by the cancer cells. uPA, encoded by the Plasminogen Activator Urokinase (PLAU) gene, have been identified to regulate the metastatic cascade by aiding degradation of surrounding ECM, thus enhancing cancer invasion *(32)*. Furthermore, previous studies have also shown that proliferation of cancer cells was higher near perfused than non-perfused vessels or at larger distances from perfused vessels *(33, 34)*.

Proximal tumors responded well to both chemo and immunotherapy. Doxorubicin, an anthracycline-based chemotherapy drug, primarily intercalates into the DNA and inhibits topoisomerase II (TOP2) in proliferating cancer cells, eventually leading to cancer cell death *(35)*. Perfusion of doxorubicin to treat proximal tumors resulted in extensive cancer cell death for high drug concentrations (100, 10 and 1 µM), which was evident from decreasing tumor volumes post-treatment. Overall, testing doxorubicin under a dynamic flow-based culture exhibited a dose dependent reduction in tumor growth with decreased doxorubicin activity below 1 µM. Induction of apoptotic pathways are generally associated with the activation of a group of cysteine proteases, called caspases *(36)*. Specifically, Caspase 3 catalyzes cleavage of key cellular proteins, DNA fragmentation, cell rupture and formation of apoptotic bodies *(37)*. Caspase 3/7 activity, found significantly enhanced in 100-1 µM doxorubicin treated cultures, resulted in extensive cell death and subsequent reduction in tumor volumes. Doxorubicin uptake and colocalization in cellular nuclei decreased below 1 µM concentrations, which further indicated decreased uptake and drug resistance in tumors. The results of the proteome array also suggested chemo-resistance in tumors below 1 µM, under the dynamic flow-based culture. Comparison of the protein expression profiles of both treated groups (1 and 0.1 µM) with the non-treated group, showed a substantial increase in the expression of several cytokines, chemokines, and growth factors, post-chemotherapy. Proteins such as decorin, enolase-2, foxC2, galectin-3, IL-6, lumican, CCL2/MCP-1, M-CSF, GM-CSF, Osteopontin (OPN), uPA, and VEGF were all significantly higher in expression post chemotherapy. These cytokines and growth factors secreted by cancer cells play a major role in cancer cell proliferation and survival, and progression as well as formation of tumor stroma *(38)*. Thus, overexpression of these proteins could provide proliferative and anti-apoptotic signals aiding tumor survival and helping the tumor escape drug-mediated apoptosis. Cytokines and chemokines, such as interleukin-6 (IL-6), interleukin-8 (IL-8), monocyte chemoattractant protein 1 (MCP-1), monocyte colony stimulating factor (M-CSF), and granulocyte-macrophage colony stimulating factor (GM-CSF), are all produced by cancer cells and heavily involved in a myriad of paracrine and autocrine functions *(39)*. IL-6 and IL-8 are both known to stimulate tumor cell proliferation as well as promote angiogenesis *(40, 41)*. GM-CSF and M-CSF are both immune modulatory cytokines secreted by activated immune (macrophage, monocyte) cells, stromal (fibroblast, vascular endothelial cells) or even cancer cells, including, MDA-MB-231 cells, in response to various stimuli *(42, 43)*. Studies have also implicated GM-CSF and M-CSF in promoting tumor growth and progression *(42)*. Increased expression of FoxC2 increases the promoter activity of ABC transporters, which are known to be associated with multidrug resistance (MDR) and epithelial–mesenchymal transition (EMT) *(44, 45)*. Increased expression of Decorin, Galectin-3 and Enolase-2 post doxorubicin treatment could also potentially be used as markers of chemotherapy efficacy *(46, 47)*. All of these proteins are involved in matrix remodeling *(48)*, cell migration *(49)*, cell metabolism *(50)* and their increased expression after the doxorubicin treatment exhibits the propensity of metastatic cancer cells to evade apoptosis. Increased expression of all these factors enables the cancer cells to evade drug-induced death *(51, 52)*.

Cellular engineering of immune cells as CAR-T cells has shown great therapeutic promise against hematological tumors *(53)* but is not yet effective against solid tumors. An immune-suppressive microenvironment composed of a dense mass of tumor and stromal cells and abnormal vasculature restricts immune cell infiltration and thus suppresses immune function *(54)*. In our study, we were able to recapitulate a physiologically-relevant tumor microenvironment consisting of rapidly dividing tumor cells, stromal cells as well as extensive vasculature through and around the tumor. Thus, perfusing HER2 and CD19 targeted CAR T-cells through the central vasculature resulted in extensive CAR-T cell adhesion to vasculature wall as well as CAR-T cell infiltration into the surrounding matrix. This phenomenon could be attributed to the presence of E- and P-selectins, which enable rolling of CAR-T cells on the endothelium *(55)*. Furthermore, activated T cells express ligands, which enable them to bind to intercellular adhesion molecule 1 (ICAM-1) and vascular cell adhesion molecule 1 (VCAM-1) present on endothelial cells *(56)*. Increased expression of IFNγ post culture with anti-HER2 CAR-T cell versus B cell lymphoma specific *(57)* anti-CD19 CAR-T cells, which served as controls, revealed antigen specific T cell activation leading to antitumor activity *(58)*. Higher expression of IFNγ along with cytotoxic granzymes, which are known to mediate cellular apoptosis by diffusing through perforin pores on the plasma membrane of target cells, possibly resulted in an antiproliferative, and pro-apoptotic tumor microenvironment in HER2 targeted CAR treatments *(59, 60)*. Furthermore, higher secretion of cytokines GM-CSF and IL-13 in HER2 targeted CAR-T cell treated groups as compared to anti-CD19 CARs, also substantiated HER2 specific CAR-T cell activation. Chemokines, such as monocyte chemoattractant protein-1 (CCL2/MCP-1), and CXCL10 (interferon-γ-inducible protein 10, previously called IP-10), and monokine induced by gamma (MIG/CXCL9), play a crucial role in immune cell stimulation and immune cell recruitment to intra-tumoral sites *(61)*. Their secretion is known to be induced by IFNγ are produced by a wide range of cell types, including endothelial cells, fibroblasts, monocytes, and neutrophils *(62),(63),(64)*. Higher expression of these factors after HER2 targeted CAR-T cell treatment as compared to CD19 CARs, further validated the 3D bioprinted perfusable tumor model as an effective platform for not only immunotherapy screening but further eliciting immune responses post-treatment.

## Conclusions

In summary, our study elucidated the development of a 3D vascularized dynamic-flow based breast tumor microenvironment for chemo and immunotherapeutic applications. Controlling the spatial location of tumors substantiated that distance of tumors from a perfused vasculature affects tumor angiogenesis and cancer invasion, two of the major hallmarks of cancer. Dose dependent reduction in tumor volumes as well as drug resistance beyond 0.1 µM of doxorubicin validated the responsiveness of tumor microenvironment against therapeutics. Additionally, perfusing HER2 targeted CAR-T cells through the tumor vasculature, resulted in extensive T cell recruitment to the endothelium, suggestive of an inflammatory microenvironment. Perfusing HER2 targeted CARs through vasculature induced secretion of inflammatory cytokines and chemokines, activating CARs to generate antitumor response, which resulted in decreased tumor growth. Overall, fabrication of a physiologically-relevant in vitro 3D tumor model is not only essential for understanding critical cancer-stroma crosstalk but also identifying novel therapeutic targets and developing targeted therapies against cancer.

## Materials and Methods

### Study Design

The objective of the study was to develop a physiologically-relevant 3D tumor model for drug testing and immunotherapy purposes, employing aspiration-assisted bioprinting. A collagen/fibrin based natural composite hydrogel was used as the bioprinting substrate. Heterotypic tumor spheroids were aspirated and bioprinted at defined locations in the hydrogel substrate. These bioprinted devices were connected to perfusion, which induced tumor angiogenesis and invasion of cancer cells in the perfused endothelium. Perfusion devices were then subjected to chemotherapeutic and immunotherapeutic drug testing. Doxorubicin, a chemotherapeutic drug, was perfused through the devices for 72 h, following which, tumor volumes as well as cytokines released after drug treatment were measured. Similarly, anti HER2 CAR T-cells were perfused for 24 or 72 h through the devices and cytokines released after treatment were identified to understand the immune-cancer interactions in the tumor microenvironment.

### Cells and reagents

GFP^+^ MDA-MB-231 breast cancer cells were donated by Dr. Danny Welch, from University of Kansas (Kansas City, KS). They were cultured in Dulbecco’s Modified Eagle’s Medium (DMEM, Corning Cellgro, Manassas, VA) supplemented with 5% fetal bovine serum (FBS) (Life Technologies, Grand Island, NY), 1 mM Glutamine (Life Technologies, Carlsbad, CA) 1 mM penicillin-streptomycin (Life Technologies, Carlsbad, CA). HUVECs and HDFs were procured from Lonza (Walkersville, MD). HUVECs were cultured in MCDB 131 media (Corning, NY) supplemented with 10% FBS, 1 mM Glutamine, 1 mM penicillin-streptomycin, 0.5 mM bovine brain extract (BBE, Lonza, Walkersville, MD), 1200 U/ml heparin (Sigma-Aldrich, St. Louis, MO) and 0.25 mM endothelial cell growth supplement (ECGS, Sigma-Aldrich). HUVECs were used at passages 2 through 7. HDFs were cultured in DMEM supplemented with 10% FBS, 1% glutamine, 1% sodium pyruvate, and 1% penicillin-streptomycin. HDFs were used at passages 2 through 8. Cells were maintained at 37 °C with 5% CO_2_ in an air-humidified atmosphere. Cell culture medium was changed every 2-3 days. Sub-confluent cultures were detached from cell culture flasks using a 0.25% trypsin-0.1% EDTA solution (Life Technologies) and split to maintain cell growth. HUVECs were further transduced with tdTomato lentiviral vector to ease visualization for all experiments as described in Supporting Information.

### Tumor spheroid formation for 3D bioprinting

Tumor (H231F) spheroids comprised of 7000 HUVECs, 1000 MDA-MB-231 cells and 1000 HDFs. Cells were individually trypsinized and combined according to the above-mentioned ratio and seeded in a 96-well U-bottom cell repellant plate (Greiner Bio-One, Monroe, NC) in 75 µl of EGM-2MV media per well. Cells were cultured in the U-bottom plate for 24 h to form spheroids.

### Immunostaining of tumor spheroids

Tumor spheroids were fixed in 4% paraformaldehyde (Sigma-Aldrich) and rinsed in Dulbecco’s phosphate buffered saline (DPBS, 1X). The spheroids were permeabilized using 0.1% Triton-X100 (Sigma Aldrich, Burlington, MA) for 15 min and blocked with 10% normal goat serum (NGS, Thermo Fisher, Waltham, MA), 0.3 M glycine (Sigma-Aldrich), 0.1% Tween-20 (Sigma-Aldrich), and 1% BSA (Sigma Aldrich) in DPBS for 1 h. Samples were then incubated with rabbit monoclonal anti-PDGFR alpha antibody (1:200, Abcam, Waltham, MA) in the same blocking solution overnight at 4 °C. Afterwards, they were washed twice with DPBS and incubated with goat anti-rabbit IgG (H + L)-Alexa Fluor 647 (1:1000, Invitrogen, Waltham, MA) for 1 h at room temperature, followed by incubation with Hoechst 33258 (1:200, Sigma Aldrich) for 30 min to visualize the cell nuclei. Images were taken using a Leica SP8 DIVE Multiphoton Microscope (Leica Microsystems, Germany).

### Hydrogel preparation for 3D perfusable devices

Type I collagen was extracted from rat tails as described in the Supplementary information. Fibrinogen and thrombin were purchased from Sigma Aldrich (Burlington, MA). Equal volumes of 4 mg/ml type I collagen and 6 mg/ml fibrin *(20)* was mixed together to obtain a final concentration of 2 mg/ml collagen and 3 mg/ml fibrin. This composite hydrogel will be referred to as ‘C2F3’. Briefly, 5 µL DPBS (10X), 0.51 µL sodium hydroxide (1N NaOH), 22.4 µL EGM-2MV media, 50 µL fibrinogen (6 mg/ml), 22.2 µL collagen (9 mg/ml) and 1.5 U/ml thrombin (50 U/ml) were mixed together in this specific order. All of the above-mentioned solutions, except fibrinogen, were kept on ice prior to mixing. HDFs at a concentration of 0.5 million/ml were suspended in C2F3 and cast in the device prior to 3D bioprinting.

### Rheological studies on collagen-fibrin composite hydrogel

Rheological properties of C2F3 was characterized using a rheometer (MCR 302, Anton Paar, Austria). All rheological measurements were performed in triplicates with a 50 mm cone plate with a 1-degree gap angle at room temperature (22 °C) and 37 °C controlled by a Peltier system. The time sweep test was carried out in an angular frequency range of 1 Hz. To monitor changes in rheological properties of C2F3 incubated at two different temperatures (22 °C to mimic bioprinting condition and 37 °C to mimic incubation conditions post-bioprinting), a time sweep test was carried out under constant dynamic-mechanical conditions at a fixed angular frequency of 1 Hz and a fixed shear strain of 0.1%, which was within the linear viscosity region.

### Measurement of mechanical strength of tumor spheroids

A pulled micropipette with a final inner diameter of 70-85 µm, connected to vacuum, was used to measure the mechanical strength of MDA-MB-231, H231, and H231F spheroids. The micropipette was placed on an optical microscope (Motic, Schertz, TX) and the testing was performed in medium with a fixed aspiration pressure of 4 kPa to apply the force to spheroids for 10 min. The aspiration pressure was controlled by a pressure controller (Ultimus™ I, Nordson EFD, Westlake, OH). Videos were recorded for 10 min and images were taken every 1 min. The deformation of spheroids was calculated based on the change in length of aspiration in the micropipette tip from pre-deformed spheroids. The elastic modulus was calculated using a previously established equation, (Eq. 1) under the assumption of the homogeneous half-space model *(65, 66)*.,

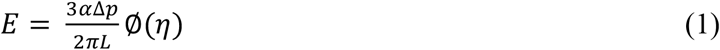

Where E is the elastic modulus, *α* is the inner radius of the micropipette, Δ*p* is the aspiration pressure, and ∅(*η*) is the geometry of pulled micropipette.

### Device fabrication and aspiration-assisted bioprinting of tumor spheroids

A poly-lactic acid (PLA)-based 3D printed structure was used as the perfusable chamber (Fig. 1A1). This structure was designed using Autodesk Inventor (Autodesk, San Rafael, CA) and then 3D printed by Ultimaker 2 (Ultimaker, Germany). This 3D printed device was coated with 1 mg/ml of poly-D lysine hydrobromide (Sigma Aldrich) overnight. Devices were then thoroughly rinsed with sterile DI water to remove excess lysine and air dried prior to printing. H231F tumor spheroids were suspended in 3 mg/ml fibrinogen prior to bioprinting. Tumor spheroids were individually aspirated from fibrinogen using a 30G blunt nozzle, having a diameter of ∼150 µm, and vacuum pressure of ∼20 mmHg. They were then bioprinted into the semi-crosslinked C2F3 matrix at a speed of 5 mm/s. The first tumor was bioprinted within 60 sec of C2F3 deposition. A total of six spheroids were bioprinted in C2F3 within 5 min. Spheroids were bioprinted at controlled distances of ∼100 µm (proximal) and ∼500 µm (distal) from the stainless-steel wire (later referred to as the ‘central vasculature’) as shown in Fig. 1. Bioprinting accuracy was measured according to the protocol described in Supplementary Information. After device fabrication, it was transferred to a humidified incubator at 37 °C and 5% CO_2_ flow for complete hydrogel crosslinking. The stainless-steel wire was removed from the device leaving behind an open channel. The open channel was flushed with cell culture media a few times. HUVECs were injected into the channel, and the device was turned every 30 min to ensure homogenous cell attachment on both upper and lower channel walls. After 60 min, the device was submerged in EGM-2MV media and incubated at 37 °C and 5% CO_2_ flow, overnight, to allow HUVECs to completely adhere to the channel walls.

### Measuring tumor viability before and after bioprinting

To analyze the effect of aspiration-assisted bioprinting on tumor viability, six tumor spheroids were bioprinted into C2F3. Post bioprinting, a viability assay was performed using a standard Cell Counting Kit-8 (CCK8) protocol (Dojindo, Rockville, MD). Briefly, CCK8 solution was diluted in culture media at a ratio of 1:10. Bioprinted samples were incubated with ∼ 350 µL of this diluted CCK8 solution for 4 h. To compare any changes in viability due to bioprinting, control samples were made by manual encapsulation of the same number of tumor spheroids in C2F3. After incubating the samples with CCK8 solution for 4 h, the culture media was collected and any changes in absorbance of the CCK8 solution was measured at 450 nm using a microplate reader (PowerWaveX, Biotek, Winooski, VT). Viability of the bioprinted tumor spheroids were normalized based on the manually-made control group. All experiments were performed in triplicates. Similarly, to analyze the effect of bioprinting on tumor structure, a circularity analysis was performed as described in the Supplementary Information.

### Perfusion, imaging and quantitative analysis of 3D bioprinted perfusion device

HUVECs were injected into the channel to form a uniform endothelial lining, as described under device fabrication. After HUVEC attachment to the channel walls, the device was connected to an external pump (Reglo Ismatec, MasterFlex, Radnor, PA) and perfused for a period of 6 days. The perfusion speed was maintained at 0.7 μL/min. Formation of endothelium was confirmed using dextran perfusion and immunostaining as described in Supplementary Information. 3D Bioprinted devices were fixed with 4% formaldehyde (Santa Cruz, California, USA), overnight. Devices were then washed with DPBS (1X) and imaged using a 16x immersion lens on the Leica SP8 DIVE multiphoton microscope. For imaging the entire region including the six bioprinted spheroids, an imaging workflow was set up on the Leica software interface, where Z stacks were collected from individually defined regions and then stitched together using the ‘mosaic merge’ function. Quantitative analysis on tumor angiogenesis was performed using the Angiotool software *(67)*.

### Drug study

Doxorubicin hydrochloride (Tocris Biosciences, Minneapolis, MN) was dissolved in dimethyl sulfoxide (DMSO) to prepare a stock concentration of 50 mM, aliquoted and stored at -30 ^°^C. In this study, doxorubicin concentration was diluted 10-folds each time, starting from a very high concentration of 100 µM up to a very low concentration of 0.0001 µM. The non-treated control group contained only the vehicle (DMSO) diluted in EGM-2MV media (0.1%). For free-standing spheroids cultured in static conditions, spheroids in well plates were directly exposed to doxorubicin for a period of 72 h. Similarly, to understand the effect of doxorubicin on spheroids encapsulated in C2F3, spheroids were first cultured in C2F3 for a period of 3 days. Then, doxorubicin, diluted in culture media (100 - 0.0001 µM), was added to these cultures and incubated for 72 h. As the MDA-MB-231 cells used were GFP^+^, images were captured daily using a 4x lens on the EVOS fluorescent microscope to assess the changes in fluorescence intensity of the cancer cells over time.

### Viability study after drug testing

After incubating free-standing and hydrogel encapsulated tumor spheroids with doxorubicin for a 72-h culture period, Alamar Blue assay (Thermo Fisher) was performed according to the manufacturer’s protocol in order to quantify the cell viability after the drug treatment. Briefly, samples were incubated with Alamar Blue solution, which was diluted at a ratio of 1:10 in culture media, for a period of 4 h. The culture supernatant was then removed and fluorescence intensity of the resorufin produced was measured at an excitation between 530–560 and an emission at 590 nm, using a fluorescence-based microplate reader (Tecan, Morrisville, NC).

### Caspase 3/7 activity assay

Induction of apoptosis after 24 h of doxorubicin treatment was quantified by analyzing Caspase 3/7 activity using the Caspase-Glo^®^ 3/7 3D assay (Promega, Madison, WI). The assay protocol was followed from the manufacturer’s website. Briefly, the Caspase Glo^®^ 3/7 3D buffer was mixed with the Caspase Glo^®^ 3/7 3D substrate. This mixture constituted the Caspase Glo^®^ 3/7 3D Reagent, which was then added to the perfusion devices. Devices were incubated at RT for 30 min. The solution was removed from the devices and luminescence was recorded on microplate reader (Tecan, Morrisville, NC).

### Proteome array

In this study, Proteome Profiler Human XL Oncology Array (R&D Systems, Minneapolis, MN) was used to identify 84 different cancer-related antibodies. The experimental groups included a drug treated group (1 µM doxorubicin) and a non-treated group (0 µM doxorubicin). Cell culture supernatant was collected after 72 h drug perfusion, aliquoted and stored at -80 ^°^C for later use. ∼300 µL of this supernatant was used for the proteome array for both drug treated and non-treated groups, and the manufacturer’s protocol was followed for this assay. ChemiDoc MP Imaging System (Bio Rad Laboratories, Hercules, CA, USA) was used to image the protein blots and the intensity of the protein blots was quantified using ImageJ.

### Designing CAR constructs and engineering CAR-T cells

Anti-CD19 CAR and anti-HER2 CAR encoding lentiviral vectors were produced and titrated as described in Supplementary Information. CAR constructs consist of CD8 alpha signal peptide, single chain variable fragment (scFv) of a CD19 or HER2 antibody, CD8 hinge domain, CD8 transmembrane domain, 4-1BB (CD137) intracellular domain and CD3ζ domain which were designed with Snapgene and synthesized via Genscript. CD8a signal peptide, CD8 hinge, CD8 transmembrane domain, 4-1BB intracellular domain and CD3ζ domain sequences were obtained from Ensembl Gene Browser and codon optimized with SnapGene by removing the restriction enzyme recognition sites that are necessary for subsequent molecular cloning steps while preserving the amino acid sequences. Anti-CD19 and anti-HER2 scFv amino acid sequences were obtained from Addgene plasmids #79125 and #85424, respectively, reverse translated to DNA sequences and codon optimized with Snapgene 5.2.4. The constructs were then cloned into a lentiviral expression vector with a multiple cloning site separated from GFP reporter via an Internal Ribosomal Entry Site (IRES).

CD8+ T cells were activated using anti-CD3/CD28 coated beads (Invitrogen) at a 1:2 ratio (beads:cells) and infected with anti-CD19 CAR, anti-HER2 CAR or empty lentivectors with multiplicity of infection (MOI) of 3-10. Cells were then expanded in complete RPMI 1640 medium supplemented with 10% FBS (Atlanta Biologicals, Flowery Branch, GA), 1% penicillin/streptomycin (Corning Cellgro) and 20 ng/ml of IL-2 for 10-12 days and cultured at 37 °C and 5% CO_2_ supplemented incubators. Respective viruses were added 24 h after the activation. Cells were expanded for 12-20 days and cytotoxicity assays were performed following their expansion.

### Anti-HER2 CAR-T cell perfusion and imaging

To understand the efficacy of anti HER2 CAR T-cells in killing MDA-MB-231 cells, static cultures were conducted prior to perfusion studies, as described in Supplementary Information. CAR T-cells were labelled with cell trace violet (Invitrogen, Waltham, MA) prior to using them for perfusion experiments, following manufacturer’s protocol. After staining, CAR T-cells were suspended in 1:1 mixture of RPMI and EGM-2MV media. This media combination was further supplemented with IL-2 at a 1:500 dilution. T cells, suspended in this media combination were introduced in 3D bioprinted perfusion device and perfused for 24 or 72 h. Perfusion devices were then fixed with 4% para-formaldehyde and imaged using a 16x immersion lens on SP8 multiphoton microscope (Leica Microsystems, Morrisville, NC). Tumor volumes were quantified by imaging individual tumors post bioprinting

### Supernatant analysis

Qbeads immunoassay was used for analyzing cytokines and chemokines secreted after CAR T perfusion for 24 or 72 h. Capture beads fluorescently tagged with a unique signature and coated with capture antibodies directed against a specific analyte were incubated with cell culture supernatants in a 96-well V-bottom plate. Once the analyte was bound by the capture beads, a fluorescent detection antibody was added to the reaction which then bound to the analyte already bound to the beads. To maximize analyte sensitivity and reduce fluorescence background, the bead/analyte/detection are washed. Data were acquired on Intellicyt iQue Screener Plus (Albuquerque, NM). The fluorescence signal was associated with the bead complex and the fluorescence intensity directly correlated to the quantity of bound analyte. Data were represented as mean fluorescence intensity. To assess the production of multiple secreted proteins and cytokines, including GM-CSF, Granzyme A, IFN-γ, IL-13, CCL2 (MCP1), MIG and IP10, the assay was multiplexed.

### Statistics

All data were presented as the mean ± standard deviation and analyzed by Minitab 17.3 (Minitab Inc., State College, PA) using one-way analysis of variance (ANOVA) followed by the Posthoc Tukey’s multiple comparison test. When comparing multiple groups with a single control group, a Dunnett Multiple Comparisons test was used. Statistical differences were considered significant at ^*^p < 0.05, ^**^p <0.01, ^***^p < 0.001.

## Acknowledgments

We are grateful to Dr. Danny Welch, from University of Kansas (Kansas City, KS) for providing GFP^+^ MDA-MB-231 cells. We also acknowledge the support from The Huck Institutes of Life Sciences and Materials Research Institute at Penn State (University Park, PA) and Jackson Laboratory (Farmington, CT) for providing facilities for characterization of experiments.

## Author contributions

M.D., D.U. and I.T.O developed the ideas and designed the experimental plan. M.D., M.H.K., M.N., M.D and L.K. performed the experiments. M.D. took the lead in writing the manuscript. All authors provided critical feedback and helped shape the research, analysis and manuscript, and approved the content of the manuscript. All authors contributed to writing the manuscript and agreed on the final content of the manuscript.

## Data and materials availability

All data needed to evaluate the conclusions in the paper are present in the paper and/or the Supplementary Information. Additional data related to this paper may be requested from the authors.

## References

1. J. Ferlay, M. Colombet, I. Soerjomataram, D. M. Parkin, M. Piñeros, A. Znaor, F. Bray, Cancer statistics for the year 2020: An overview. Int. J. Cancer 149, 778–789 (2021).

2. D. J. Adams, The Valley of Death in anticancer drug development: a reassessment. Trends Pharmacol. Sci. 33, 173–180 (2012).

3. M. Guo, Y. Peng, A. Gao, C. Du, J. G. Herman, Epigenetic heterogeneity in cancer. Biomark. Res. 2019 71 7, 1–19 (2019).

4. B. Gottlieb, M. Trifiro, G. Batist, Why Tumor Genetic Heterogeneity May Require Rethinking Cancer Genesis and Treatment. Trends in Cancer 7, 400–409 (2021).

5. F. S. Hodi, S. J. O’Day, D. F. McDermott, R. W. Weber, J. A. Sosman, J. B. Haanen, R. Gonzalez, C. Robert, D. Schadendorf, J. C. Hassel, W. Akerley, A. J. M. van den Eertwegh, J. Lutzky, P. Lorigan, J. M. Vaubel, G. P. Linette, D. Hogg, C. H. Ottensmeier, C. Lebbé, C. Peschel, I. Quirt, J. I. Clark, J. D. Wolchok, J. S. Weber, J. Tian, M. J. Yellin, G. M. Nichol, A. Hoos, W. J. Urba, Improved Survival with Ipilimumab in Patients with Metastatic Melanoma. N. Engl. J. Med. 363, 711–723 (2010).

6. D. B. Doroshow, M. F. Sanmamed, K. Hastings, K. Politi, D. L. Rimm, L. Chen, I. Melero, K.A. Schalper, R. S. Herbst, Immunotherapy in Non–Small Cell Lung Cancer: Facts and Hopes. Clin. Cancer Res. 25, 4592–4602 (2019).

7. P. Ghatalia, M. Zibelman, D. M. Geynisman, E. R. Plimack, Checkpoint Inhibitors for the Treatment of Renal Cell Carcinoma. Curr. Treat. Options Oncol. 18, 1–14 (2017).

8. A. Polk, I. M. Svane, M. Andersson, D. Nielsen, Checkpoint inhibitors in breast cancer – Current status. Cancer Treat. Rev. 63, 122–134 (2018).

9. F. Marofi, R. Motavalli, V. A. Safonov, L. Thangavelu, A. V. Yumashev, M. Alexander, N. Shomali, M. S. Chartrand, Y. Pathak, M. Jarahian, S. Izadi, A. Hassanzadeh, N. Shirafkan, S. Tahmasebi, F. M. Khiavi, CAR T cells in solid tumors: challenges and opportunities. Stem Cell Res. Ther. 2021 121 12, 1–16 (2021).

10. S. Toulouie, G. Johanning, Y. Shi, Chimeric antigen receptor T-cell immunotherapy in breast cancer: Development and challenges. J. Cancer 12, 1212–1219 (2021).

11. I. W. Y. Mak, N. Evaniew, M. Ghert, Lost in translation: animal models and clinical trials in cancer treatment. Am. J. Transl. Res. 6, 114 (2014).

12. Y. Nashimoto, R. Okada, S. Hanada, Y. Arima, K. Nishiyama, T. Miura, R. Yokokawa, Vascularized cancer on a chip: The effect of perfusion on growth and drug delivery of tumor spheroid. Biomaterials 229, 119547 (2020).

13. K. Haase, G. S. Offeddu, M. R. Gillrie, R. D. Kamm, K. Haase, R. D. Kamm, G. S. Offeddu, M. R. Gillrie, Endothelial Regulation of Drug Transport in a 3D Vascularized Tumor Model. Adv. Funct. Mater. 30, 2002444 (2020).

14. J. Forster, W. Harriss-Phillips, M. Douglass, E. Bezak, A review of the development of tumor vasculature and its effects on the tumor microenvironment. Hypoxia Volume 5, 21–32 (2017).

15. F. Meng, C. M. Meyer, D. Joung, D. A. Vallera, M. C. McAlpine, A. Panoskaltsis-Mortari, F. Meng, D. Joung, M. C. McAlpine, C. M. Meyer, A. Panoskaltsis-Mortari, D. A. Vallera, 3D Bioprinted In Vitro Metastatic Models via Reconstruction of Tumor Microenvironments. Adv. Mater. 31, 1806899 (2019).

16. B. Soo Kim, W.-W. Cho, G. Gao, M. Ahn, J. Kim, D.-W. Cho, B. S. Kim, W. Cho, M. Ahn, J. Kim, D. Cho, G. Gao, Construction of Tissue-Level Cancer-Vascular Model with High-Precision Position Control via In Situ 3D Cell Printing. Small Methods 5, 2100072 (2021).

17. F. Runa, S. Hamalian, K. Meade, P. Shisgal, P. C. Gray, J. A. Kelber, Tumor Microenvironment Heterogeneity: Challenges and Opportunities. Curr. Mol. Biol. Reports 2017 34 3, 218–229 (2017).

18. B. Ayan, D. N. Heo, Z. Zhang, M. Dey, A. Povilianskas, C. Drapaca, I. T. Ozbolat, Aspiration-assisted bioprinting for precise positioning of biologics. Sci. Adv. 6, eaaw5111 (2020).

19. D. N. Heo, B. Ayan, M. Dey, D. Banerjee, H. Wee, G. S. Lewis, I. T. Ozbolat, Aspiration-assisted bioprinting of co-cultured osteogenic spheroids for bone tissue engineering. Biofabrication (2021), doi:10.1088/1758-5090/abc1bf.

20. M. Dey, B. Ayan, M. Yurieva, D. Unutmaz, I. T. Ozbolat, Studying Tumor Angiogenesis and Cancer Invasion in a Three-Dimensional Vascularized Breast Cancer Micro-Environment. Adv. Biol. 2100090, 1–17 (2021).

21. T. A. Moo, R. Sanford, C. Dang, M. Morrow, Overview of Breast Cancer Therapy. PET Clin. 13, 339–354 (2018).

22. J. Tchou, Y. Zhao, B. L. Levine, P. J. Zhang, M. M. Davis, J. J. Melenhorst, I. Kulikovskaya, A. L. Brennan, X. Liu, S. F. Lacey, A. D. Posey, A. D. Williams, A. So, J. R. Conejo-Garcia, G. Plesa, R. M. Young, S. McGettigan, J. Campbell, R. H. Pierce, J. M. Matro, A. M. DeMichele, A. S. Clark, L. J. Cooper, L. M. Schuchter, R. H. Vonderheide, C. H. June, Safety and efficacy of intratumoral injections of chimeric antigen receptor (CAR) T cells in metastatic breast cancer. Cancer Immunol. Res. 5, 1152–1161 (2017).

23. K. Tamada, D. Geng, Y. Sakoda, N. Bansal, R. Srivastava, Z. Li, E. Davila, Redirecting gene-modified T cells toward various cancer types using tagged antibodies. Clin. Cancer Res. 18, 6436–6445 (2012).

24. P. Li, L. Yang, T. Li, S. Bin, B. Sun, Y. Huang, K. Yang, D. Shan, H. Gu, H. Li, The Third Generation Anti-HER2 Chimeric Antigen Receptor Mouse T Cells Alone or Together With Anti-PD1 Antibody Inhibits the Growth of Mouse Breast Tumor Cells Expressing HER2 in vitro and in Immune Competent Mice. Front. Oncol. 10, 1143 (2020).

25. S. Ghaffari, R. L. Leask, E. A. V. Jones, Flow dynamics control the location of sprouting and direct elongation during developmental angiogenesis. Dev. 142, 4151–4157 (2015).

26. R. Kalluri, M. Zeisberg, Fibroblasts in cancer. Nat. Rev. Cancer 2006 65 6, 392–401 (2006).

27. E. D’Arcangelo, N. C. Wu, J. L. Cadavid, A. P. McGuigan, The life cycle of cancer-associated fibroblasts within the tumour stroma and its importance in disease outcome. Br. J. Cancer 2020 1227 122, 931–942 (2020).

28. J. A. Nagy, S.-H. Chang, A. M. Dvorak, H. F. Dvorak, Why are tumour blood vessels abnormal and why is it important to know? Br. J. Cancer 100, 865–9 (2009).

29. P. A. Galie, D. H. T. Nguyen, C. K. Choi, D. M. Cohen, P. A. Janmey, C. S. Chen, Fluid shear stress threshold regulates angiogenic sprouting. Proc. Natl. Acad. Sci. U. S. A. 111, 7968–7973 (2014).

30. D. Hanahan, R. A. Weinberg, Hallmarks of cancer: The next generation Cell 144, 646–674 (2011).

31. S. Crotti, M. Piccoli, F. Rizzolio, A. Giordano, D. Nitti, M. Agostini, Extracellular Matrix and Colorectal Cancer: How Surrounding Microenvironment Affects Cancer Cell Behavior? J. Cell. Physiol. 232, 967–975 (2017).

32. C. M. Novak, E. N. Horst, C. C. Taylor, C. Z. Liu, G. Mehta, Fluid shear stress stimulates breast cancer cells to display invasive and chemoresistant phenotypes while upregulating PLAU in a 3D bioreactor. Biotechnol. Bioeng. 116, 3084–3097 (2019).

33. J. Bussink, J. H. A. M. Kaanders, P. F. J. W. Rijken, C. A. Martindale, A. J. Van Der Kogel, Multiparameter analysis of vasculature, perfusion and proliferation in human tumour xenografts. Br. J. Cancer 1998 771 77, 57–64 (1998).

34. J. Forster, W. Harriss-Phillips, M. Douglass, E. Bezak, A review of the development of tumor vasculature and its effects on the tumor microenvironment. Hypoxia (2017), doi:10.2147/hp.s133231.

35. A. Bodley, L. F. Liu, M. Israel, R. Seshadri, Y. Koseki, F. C. Giuliani, S. Kirschenbaum, R. Silber, M. Potmesil, DNA Topoisomerase II-mediated Interaction of Doxorubicin and Daunorubicin Congeners with DNA. Cancer Res. 49 (1989).

36. T. J. Fan, L. H. Han, R. S. Cong, J. Liang, Caspase Family Proteases and Apoptosis. Acta Biochim. Biophys. Sin. (Shanghai). 37, 719–727 (2005).

37. A. G. Porter, R. U. Jänicke, Emerging roles of caspase-3 in apoptosis. Cell Death Differ. 1999 62 6, 99–104 (1999).

38. V. Levina, Y. Su, B. Nolen, X. Liu, Y. Gordin, M. Lee, A. Lokshin, E. Gorelik, Chemotherapeutic drugs and human tumor cells cytokine network. Int. J. Cancer 123, 2031–2040 (2008).

39. B. Dankbar, T. Padró, R. Leo, B. Feldmann, M. Kropff, R. M. Mesters, H. Serve, W. E. Berdel, J. Kienast, Vascular endothelial growth factor and interleukin-6 in paracrine tumor-stromal cell interactions in multiple myeloma. Blood 95, 2630–2636 (2000).

40. M. B. Nilsson, R. R. Langley, I. J. Fidler, Interleukin-6, Secreted by Human Ovarian Carcinoma Cells, Is a Potent Proangiogenic Cytokine. Cancer Res. 65, 10794–10800 (2005).

41. R. M. Strieter, M. D. Burdick, J. Mestas, B. Gomperts, M. P. Keane, J. A. Belperio, Cancer CXC chemokine networks and tumour angiogenesis. Eur. J. Cancer 42, 768–778 (2006).

42. I. S. Hong, Stimulatory versus suppressive effects of GM-CSF on tumor progression in multiple cancer types. Exp. Mol. Med. 2016 487 48, e242–e242 (2016).

43. R. Ahmad, R. Thomas, F. Al-Rashed, N. Akhter, F. Al-Mulla, ACSL1 Regulates TNFα-Induced GM-CSF Production by Breast Cancer MDA-MB-231 Cells. Biomol. 2019, Vol. 9, Page 555 9, 555 (2019).

44. M. Saxena, M. A. Stephens, H. Pathak, A. Rangarajan, Transcription factors that mediate epithelial–mesenchymal transition lead to multidrug resistance by upregulating ABC transporters. Cell Death Dis. 2011 27 2, e179–e179 (2011).

45. Y. Chen, G. Deng, Y. Fu, Y. Han, C. Guo, L. Yin, C. Cai, H. Shen, S. Wu, S. Zeng, FOXC2 Promotes Oxaliplatin Resistance by Inducing Epithelial-Mesenchymal Transition via MAPK/ERK Signaling in Colorectal Cancer. Onco. Targets. Ther. 13, 1625 (2020).

46. A. Shafiq, J. Moore, A. Suleman, S. Faiz, O. Farooq, A. Arshad, M. Tehseen, A. Zafar, S. H. Ali, N. U. Din, A. Loya, N. Siddiqui, F. K. Rehman, Elevated Soluble Galectin-3 as a Marker of Chemotherapy Efficacy in Breast Cancer Patients: A Prospective Study. Int. J. Breast Cancer 2020 (2020), doi:10.1155/2020/4824813.

47. E. Henke, R. Nandigama, S. Ergün, Extracellular Matrix in the Tumor Microenvironment and Its Impact on Cancer Therapy. Front. Mol. Biosci. 6, 160 (2020).

48. T. Neill, L. Schaefer, R. V. Iozzo, Decorin as a multivalent therapeutic agent against cancer. Adv. Drug Deliv. Rev. 97, 174–185 (2016).

49. Effects of the carbohydrate-binding protein galectin-3 on the invasiveness of human breast carcinoma cells - Le Marer - 1996 - Journal of Cellular Physiology - Wiley Online Library (available at https://onlinelibrary.wiley.com/doi/pdf/10.1002/(SICI)1097-4652(199607)168:1%3C51::AID-JCP7%3E3.0.CO;2-7?casa_token=o8cUtPs7uSwAAAAA:IHZFB863rlCaoeXv7jT6AZ4TXlMBiGSDvxsohNd7UbOTQX2jytOMSwmTBX91lCyn0ecM64lCL8VQFZQ).

50. T. Vizin, J. Kos, Gamma-enolase: a well-known tumour marker, with a less-known role in cancer. Radiol. Oncol. 49, 217 (2015).

51. C. Ding Caijun Wu, X. Tan, X. Hu, M. Zhou, Myeloid Cells high Differentiation of Immunosuppressive Ly6C Gemcitabine Treatment Favors Tumor Microenvironment following. J Immunol Ref. 204, 212–223 (2021).

52. M. Zhang, Q. Liu, L. Li, J. Ning, J. Tu, X. Lei, Z. Mo, S. Tang, Cytoplasmic M-CSF facilitates apoptosis resistance by inhibiting the HIF - 1α/BNIP3/Bax signalling pathway in MCF-7 cells. Oncol. Rep. 41, 1807–1816 (2019).

53. Z. Zhao, Y. Chen, N. M. Francisco, Y. Zhang, M. Wu, The application of CAR-T cell therapy in hematological malignancies: advantages and challenges. Acta Pharm. Sin. B 8, 539–551 (2018).

54. L. Springuel, C. Lonez, B. Alexandre, E. Van Cutsem, J. P. H. Machiels, M. Van Den Eynde, H. Prenen, A. Hendlisz, L. Shaza, J. Carrasco, J. L. Canon, M. Opyrchal, K. Odunsi, S. Rottey, D. E. Gilham, A. Flament, F. F. Lehmann, Chimeric Antigen Receptor-T Cells for Targeting Solid Tumors: Current Challenges and Existing Strategies. BioDrugs 33, 515–537 (2019).

55. J. C. Nolz, G. R. Starbeck-Miller, J. T. Harty, Naive, effector and memory CD8 T-cell trafficking: parallels and distinctions. Immunotherapy 3, 1223 (2011).

56. C. Y. Slaney, M. H. Kershaw, P. K. Darcy, Trafficking of T Cells into Tumors. (2014), doi:10.1158/0008-5472.CAN-14-2458.

57. B. C. Miller, M. V. Maus, CD19-Targeted CAR T Cells: A New Tool in the Fight against B Cell Malignancies. Oncol. Res. Treat. 38, 683–690 (2015).

58. D. Alizadeh, R. A. Wong, S. Gholamin, M. Maker, M. Aftabizadeh, X. Yang, J. R. Pecoraro, J. D. Jeppson, D. Wang, B. Aguilar, R. Starr, C. B. Larmonier, N. Larmonier, M. H. Chen, X. Wu, A. Ribas, B. Badie, S. J. Forman, C. E. Brown, IFNγ is critical for CAR T cell–mediated myeloid activation and induction of endogenous immunity. Cancer Discov. 11, 2248–2265 (2021).

59. F. Castro, A. P. Cardoso, R. M. Gonçalves, K. Serre, M. J. Oliveira, Interferon-gamma at the crossroads of tumor immune surveillance or evasion. Front. Immunol. 9, 847 (2018).

60. I. Voskoboinik, J. C. Whisstock, J. A. Trapani, Perforin and granzymes: function, dysfunction and human pathology. Nat. Rev. Immunol. 2015 156 15, 388–400 (2015).

61. K. Kohli, V. G. Pillarisetty, T. S. Kim, Key chemokines direct migration of immune cells in solid tumors. Cancer Gene Ther. 2021 291 29, 10–21 (2021).

62. S. L. Deshmane, S. Kremlev, S. Amini, B. E. Sawaya, Monocyte Chemoattractant Protein-1 (MCP-1): An Overview. https://home.liebertpub.com/jir 29, 313–325 (2010).

63. M. Liu, S. Guo, J. K. Stiles, The emerging role of CXCL10 in cancer. Oncol. Lett. 2, 583–589 (2011).

64. R. Tokunaga, W. Zhang, M. Naseem, A. Puccini, M. D. Berger, S. Soni, M. McSkane, H. Baba, H. J. Lenz, CXCL9, CXCL10, CXCL11/CXCR3 axis for immune activation – A target for novel cancer therapy. Cancer Treat. Rev. 63, 40–47 (2018).

65. M. Sato, D. P. Theret, L. T. Wheeler, N. Ohshima, R. M. Nerem, Application of the Micropipette Technique to the Measurement of Cultured Porcine Aortic Endothelial Cell Viscoelastic Properties. (1990) (available at https://biomechanical.asmedigitalcollection.asme.org).

66. D. J. Richards, Y. Li, C. M. Kerr, J. Yao, G. C. Beeson, R. C. Coyle, X. Chen, J. Jia, B. Damon, R. Wilson, E. Starr Hazard, G. Hardiman, D. R. Menick, C. C. Beeson, H. Yao, T. Ye, Y. Mei, Human cardiac organoids for the modelling of myocardial infarction and drug cardiotoxicity. Nat. Biomed. Eng. 2020 44 4, 446–462 (2020).

67. E. Zudaire, L. Gambardella, C. Kurcz, S. Vermeren, C. Ruhrberg, Ed. A Computational Tool for Quantitative Analysis of Vascular Networks. PLoS One 6, e27385 (2011).

