## Supplementary Information for "3D Bioprinted perfusable and vascularized breast tumor model for dynamic screening of chemotherapeutics and CAR-T cells"

### Supplementary Materials and Methods

#### Transduction of HUVECs

HUVECs (Lonza, Walkersville, MD) were transduced at passage 2 (~50% confluency) with EF1 tdTomato lentiviral vector (Vectalys, Toulouse, France). A multiplicity of infection (M.O.I) of 20 was maintained for the transduction process. Briefly, a transduction mix was prepared by adding a measured amount of viral vector solution in complete culture media and 800 µg/ml polybrene (Sigma). This transduction mix was then transferred to a cell flask at ~50% confluency and incubated for 8 h. The transduction mix was then discarded and the flask was rinsed with DPBS(1X) and replenished with culture media. Cells were then allowed to grow in flasks for another 48 h before sorting them on a MoFlo Astrios sorter (Beckman Coulter, Pasadena, CA) for the brightest cells. The brightest cells were then collected for further cell culture.

#### Collagen Extraction

Collagen type-I was extracted from rat tails according to a previously published protocol *(1)*. Collagen fibers were extracted from rat tail tendons, dissolved in 0.02 M acetic acid (Sigma Aldrich) and then subsequently freeze-dried to obtain collagen sponges. These sponges were again re-dissolved in acetic acid at a desired concentration, centrifuged to remove insoluble impurities, and then sterilized using Spectra/Por 1 dialysis tubing (6–8 kDa MWCO) (Spectrum Labs, Rancho Dominguez, CA).

**Ultrastructural analysis of the tumor spheroids and hydrogel**

Field emission scanning electron microscopy (SEM, Apreo, Thermo Fisher Scientific, Waltham, MA) was used to investigate the hydrogel architecture as well as to assess surface topography of tumor spheroids. Tumor spheroids were harvested after 24 h of culture in U-bottom well plates. Hydrogel samples comprised of 2 mg/ml collagen (C2), 3 mg/ml fibrin (F3) and the C2F3 composite hydrogel. All samples were fixed in 4% paraformaldehyde (Santa Cruz Biotechnology, Dallas, TX) overnight. Samples were then washed in DPBS (1X) to remove the fixative. Next, they were dehydrated using graded ethanol solutions (25, 50, 70, 90, 100%). To ensure complete removal of water, samples were further dried in a critical point dryer (CPD300, Leica EM, Wetzlar, Germany) for 4 h. On complete dehydration, they were sputter coated with iridium using a sputter coater (Leica) and observed at an accelerating voltage of 3-5 keV on the Apreos SEM (Thermo Fisher).

**Measurement of bioprinting accuracy**

To evaluate the accuracy and precision of bioprinting of H231F spheroids in C2F3, spheroids were bioprinted at a predetermined target position on a micrometer calibration ruler. The calibration ruler was placed at the bottom of a Petri dish and recorded by a microscopic camera (Plugable USB Digital Microscope, Plugable Technologies, Redmond, WA) to monitor the target position. A total of 246 spheroids were bioprinted and analyzed by ImageJ (National Institutes of Health (NIH), MD, USA). Accuracy was represented by the root mean square error (RSME) and calculated using the following equation:

$$RMSE = \sqrt{\frac{\sum_{i=1}^{n}[(X_{Target} - X_i)^2 + (Y_{Target} - Y_i)^2]}{n}} \quad (1)$$

where, $X_{Target}$ and $Y_{Target}$ are the $X$ and $Y$ coordinates of the target position, respectively, $X_i$ and $Y_i$ are the positions of bioprinted spheroid measured in $X$ and $Y$ axes, respectively, and $n$ is the sample size. Precision was represented as the square root of the standard deviation.

**Measuring tumor circularity after bioprinting**

To represent the effect of bioprinting on the deformation of spheroids, H231F spheroids ($n$ = 50) were bioprinted in C2F3 matrix. Free-standing spheroids in 96-well plates before bioprinting were used as a control group. To quantify the morphology of spheroids, circularity was calculated using ImageJ software according to the following equation:

$$C = 4\pi \times \frac{A}{P^2} \quad (2)$$

where $C$ represents circularity, $A$ is the area, and $P$ is the perimeter of the bioprinted spheroid. The value "0" indicated an infinitely elongated polygon and "1" indicated a perfectly circular shape.

**Dextran perfusion for diffusional permeability**

Diffusional permeability was measured for the engineered vasculature by perfusing 20 µg/ml (Fluorescein isothiocyanate) FITC-conjugated 70 kDa dextran (Sigma Aldrich) in EGM-2MV media, for 40 min. Images were captured every 5 minutes using a fluorescence microscope (AxioObserver, Zeiss, NY). The diffusion of dextran and subsequent change in fluorescence intensity was measured using ImageJ. The diffusional permeability was then quantified based on the following equation:

$$P_d = \frac{1}{I_1 - I_b}\left(\frac{I_2 - I_1}{t}\right)\frac{d}{4} \quad (3)$$

Measurements were all performed on channels with and without endothelium ($n$=3).

**Lentiviral production and titration**

Cloned lentiviral constructs including anti-CD19 CAR and anti-HER2 CAR encoding vectors were co-transfected with the packaging plasmids VSVG, pLP1 and pLP2 into 293 cells using Lipofectamine™ 3000 (Invitrogen) according to the manufacturer's protocol. Viral supernatants were collected 24 - 48 h post-transfection**,** filtered through a 0.45 μm syringe filter (Millipore) to remove cellular debris**,** and concentrated with Lenti-X (Invitrogen) according to the manufacturer's protocol. Lentivirus supernatant stocks were aliquoted and stored at −80 °C. To measure viral titers, virus preps were serially diluted on Jurkat cells. 72 h after infection, $GFP^+$ cells were counted using flow cytometry and the number of cells transduced with virus supernatant was calculated as infectious units/per mL. The cells were cultured in complete RPMI 1640 medium (RPMI 1640 supplemented with 10% FBS; Atlanta Biologicals, Lawrenceville, GA), 8% GlutaMAX (Life Technologies), 8% sodium pyruvate, 8% MEM vitamins, 8% MEM nonessential amino acid, and 1% penicillin/streptomycin (all from Corning Cellgro) for 72 h. %0.05 Trypsin-0.53 mM EDTA (Corning Cellgro) was used to detach adherent cells.

**Anti HER2 CAR-T cell static culture**

Static cultures were conducted with aHER2 CAR-T cells using two types of tumor spheroids, similar to the chemotherapy experiments. Monocellular tumor spheroids containing MDA-MB-231 cells only and co-cultured tumor spheroids (H231F), were used. CAR T to MDA-MB-231 cell ratio was varied as 10:1, 20:1, 40:1 and 80:1. Control group consisted of tumor spheroids without the presence of CAR-T cells. CAR-T cells were suspended in a 1:1 mixture of RPMI and EGM-2MV media, supplemented with IL-2 at 1:500 dilution. Tumor spheroids, cultured in 96 U-bottom well plates were then directly exposed to varying concentrations of CAR-T cells and cultured for 72 h. Fluorescent images were taken daily using the EVOS FL Auto microscope, at a constant light and exposure setting for all groups, to monitor the change in GFP intensity of the cancer cells over time. The change in GFP intensity was quantified using ImageJ.

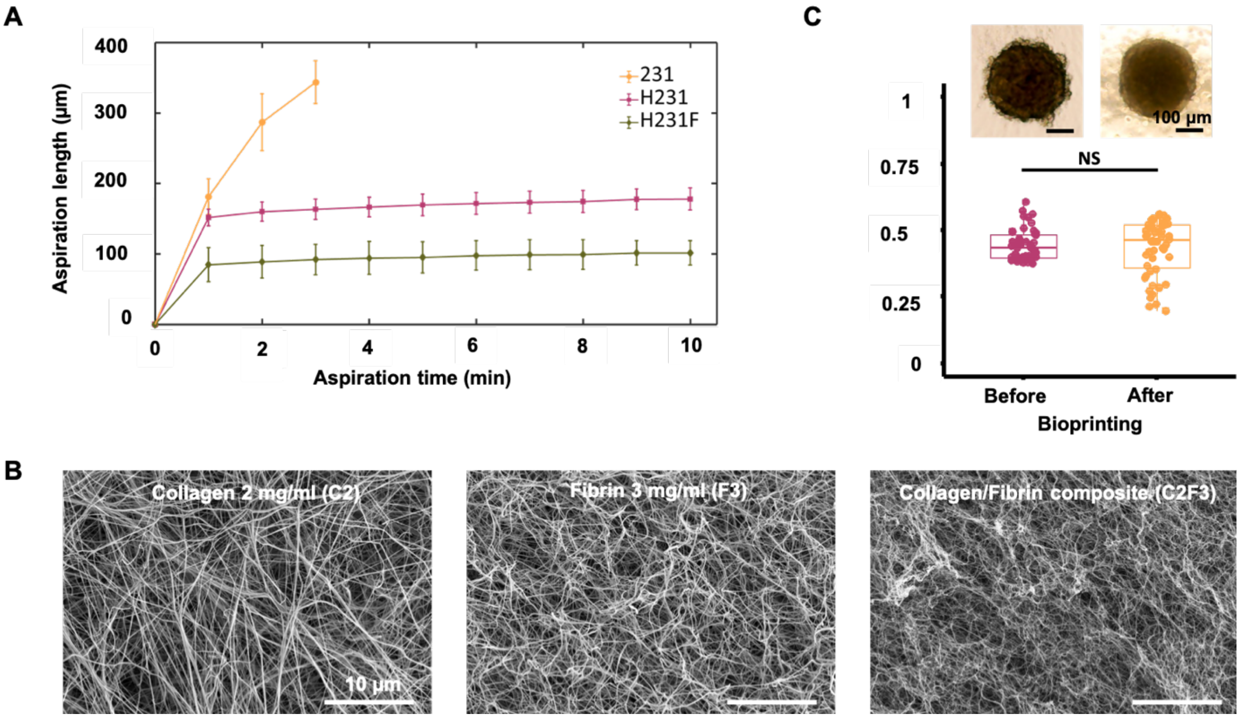


**Figure S1**: (A) Graphical representation of spheroid aspiration length over time (*n*=3). (B) SEM images of 2 mg/ml collagen (C2), 3 mg/ml fibrin (F3) and the composite hydrogel, C2F3. (C) Graphical representation of H231F tumor spheroid circularity measured before and after aspiration-assisted bioprinting (*n*=50, $p^{***} < 0.001$, $p^{**} < 0.01$, $p^{*} < 0.05$).

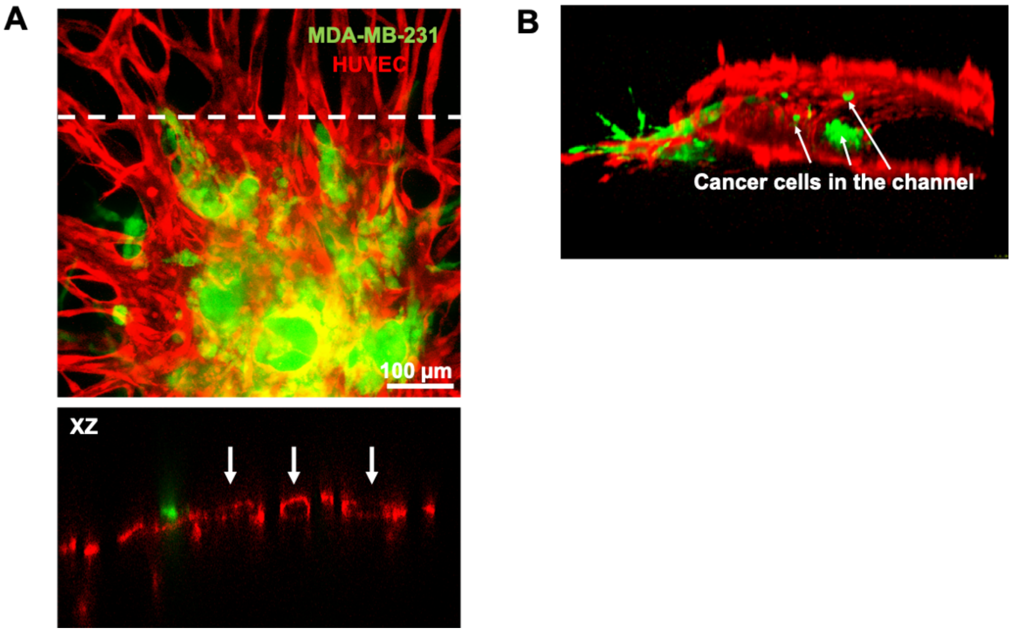


**Figure S2**: (A) An orthogonal projection across XZ plane revealing angiogenic sprouts containing hollow capillaries. White arrows are marked to represent the circular cross-section of vessels. (B) 3D Reconstruction of the perfused central vasculature containing MDA-MB-231 cells. White arrows indicate MDA-MB-231 cells trying to invade into the vasculature.

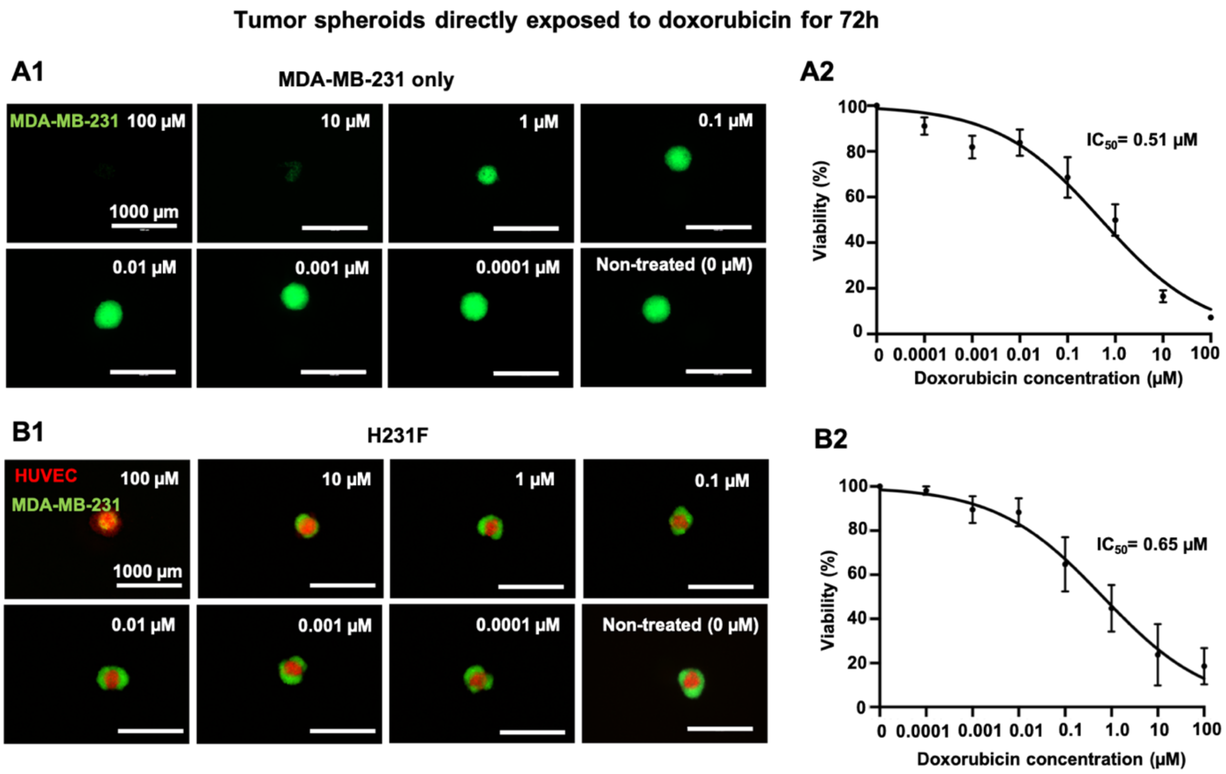


**Figure S3**: Exposure of free-standing tumor spheroids to doxorubicin. (A1) Fluorescent images of homocellular MDA-MB-231 tumor spheroids (MDA-MB-231-only) taken after 72 h of doxorubicin treatment. (A2) Dose-response curve for MDA-MB-231 spheroids treated with a range of 0 – 100 µM of doxorubicin concentration. (B1) Fluorescent images of co-cultured H231F tumor spheroids taken after 72 h of doxorubicin treatment. (B2) Dose-response curve H231F spheroids treated with a range of 0 – 100 µM doxorubicin concentration (*n*=3 for all).

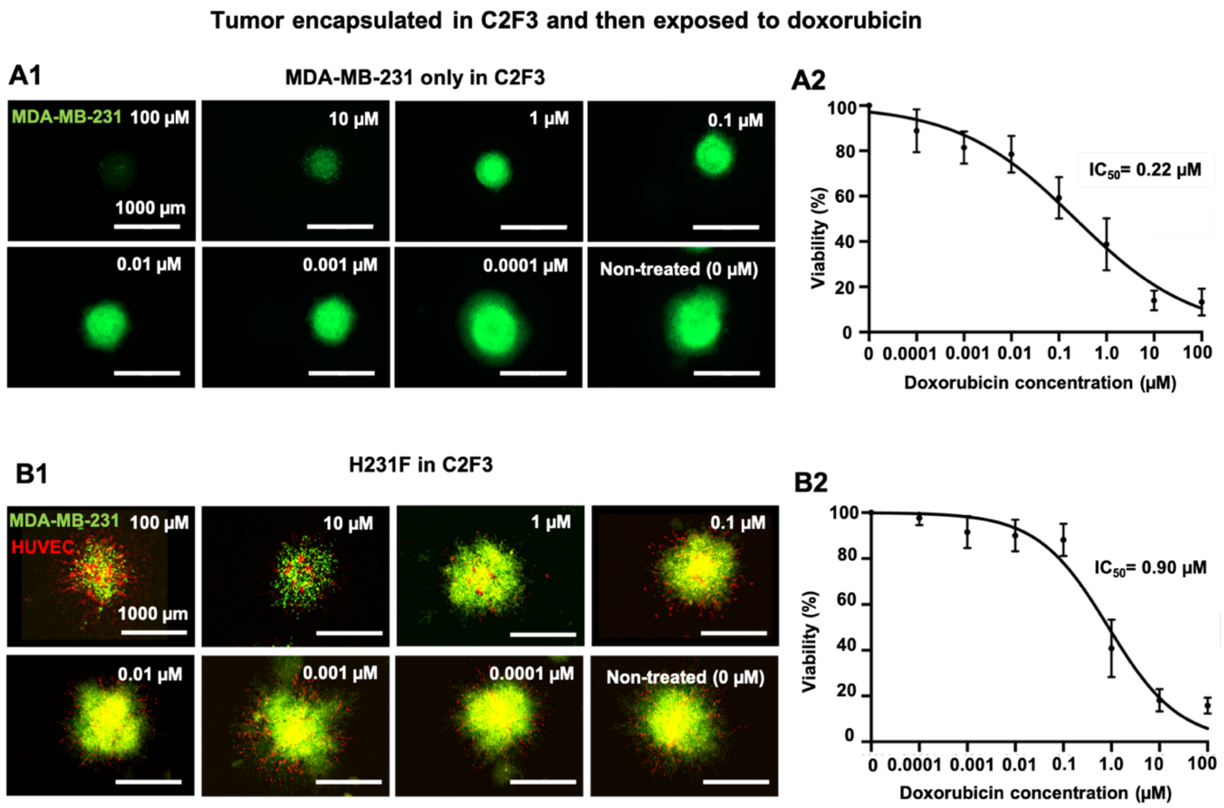


**Figure S4**: Exposure of hydrogel encapsulated tumor spheroids to doxorubicin. (A1) Fluorescent images of C2F3 encapsulated homocellular MDA-MB-231 spheroids after 72 h of doxorubicin treatment and (A2) the corresponding dose-response curve for MDA-MB-231 spheroids. (B1) Fluorescent images of C2F3 encapsulated co-cultured H231F spheroids after 72 h of doxorubicin treatment and (B2) the corresponding dose-response curve for H231F spheroids ($n$=3 for all).

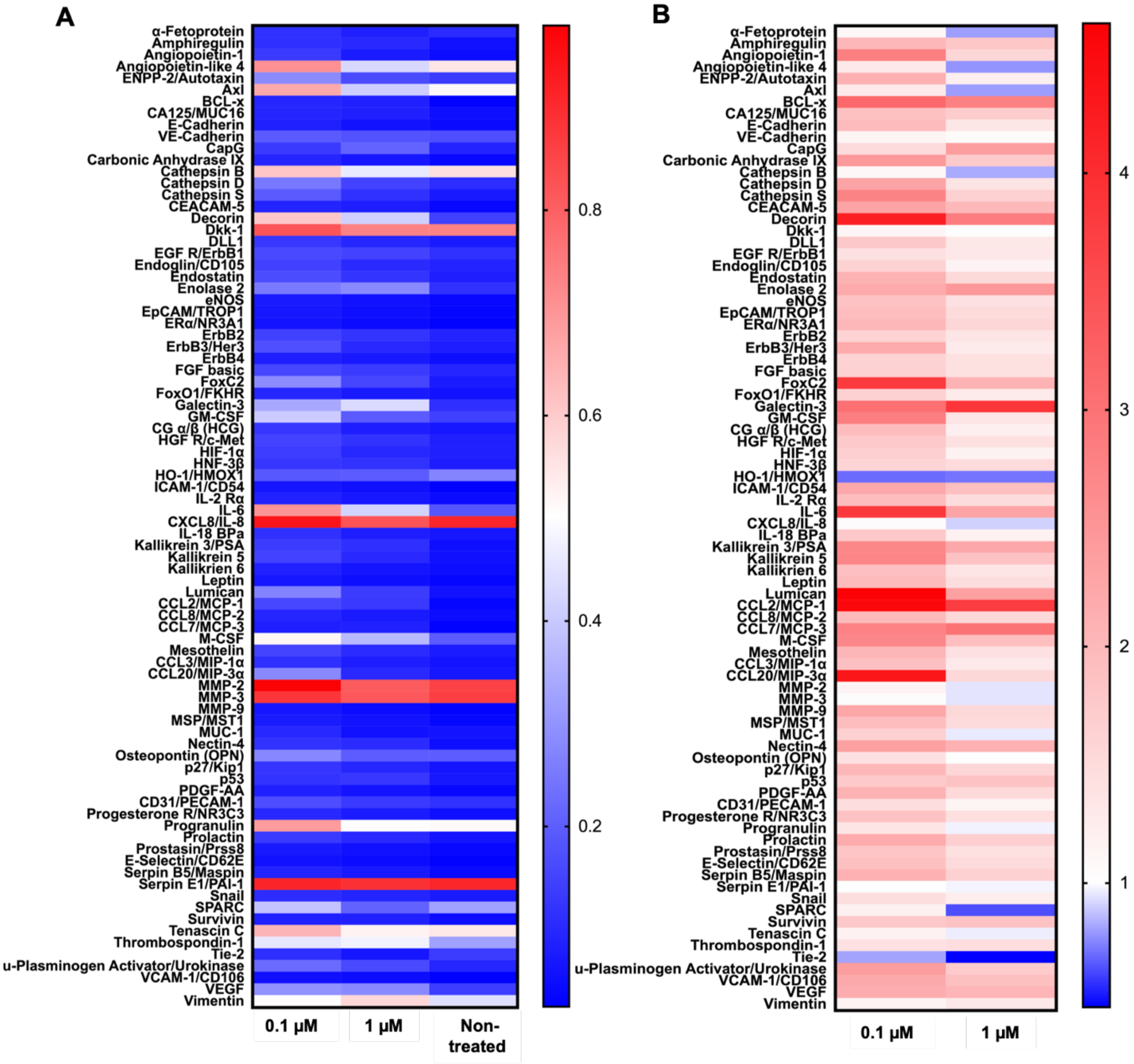


**Figure S5**: (A) Heatmap and (B) fold change in protein expression of 84 oncology-related proteins. Blue and red denoted the fold-change value lower and higher than the non-treated group, respectively. Non-treated group was normalized to 1 (*n*=3 for all).

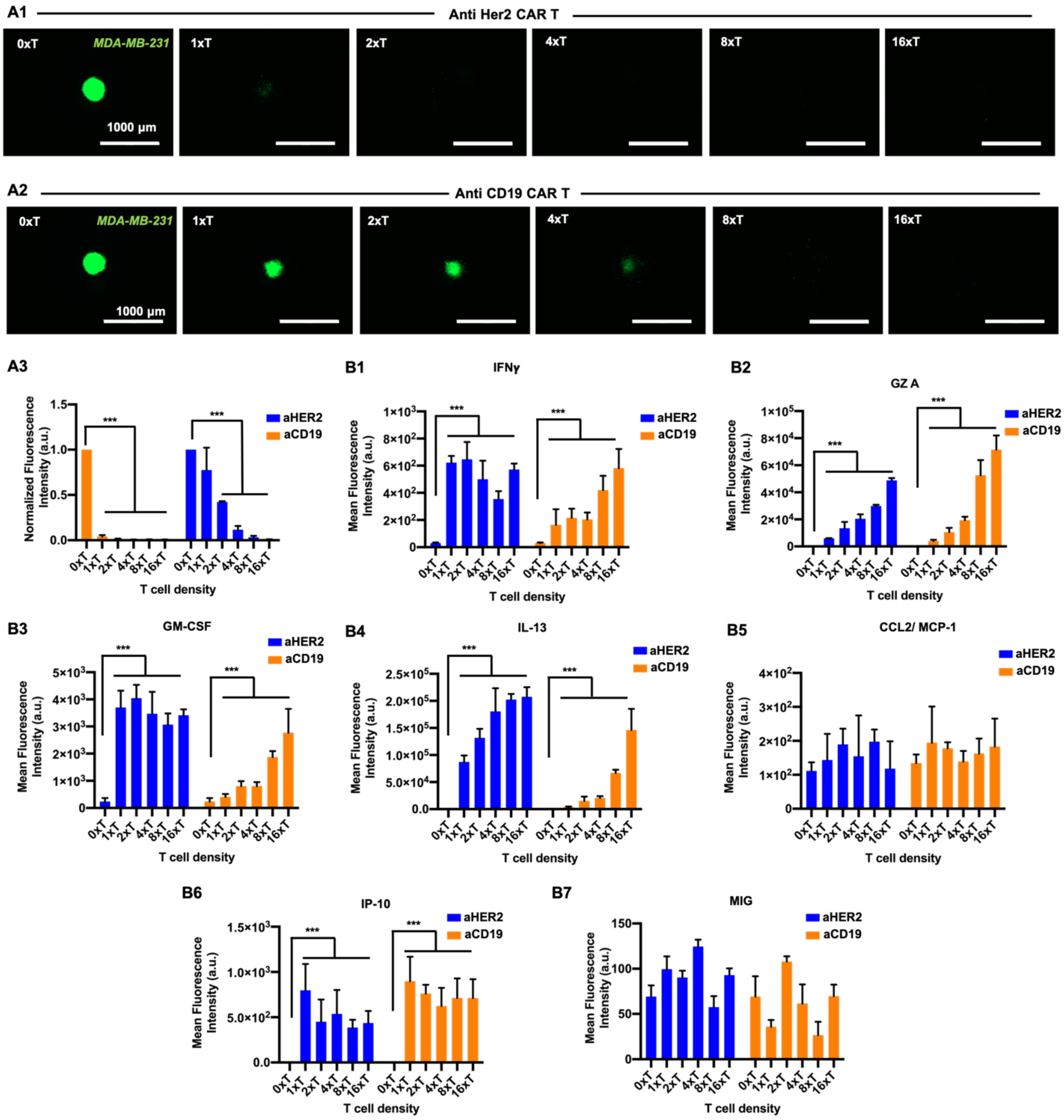


**Figure S6**: Adoptive T-cell therapy on *in vitro* static cultures of homotypic MDA-MB-231. CAR-T cell density was varied as 1xT (1:1), 2xT (2:1), 4xT (4:1), 8xT (8:1), 16xT (16:1). 0xT denotes the non-treated control culture. Fluorescent images of MDA-MB-231-only spheroids after 72 h of (A1) anti HER2 CAR-T cell and (A2) anti CD19 CAR-T cell treatment. (A3) Graphical representation of normalized fluorescence intensity of anti HER2 CAR-T cell and anti CD19 CAR-

T cell treatment. Graphical representation of the mean fluorescence intensity of the cytokines and chemokines secreted after 72 h of CAR-T cell perfusion including (B1) IFNγ, (B2) Granzyme A, (B3) GM-CSF, (B4) IL-13, (B5) CCL2/ MCP-1, (B6) CXCL10/ IP-10, and (B7) MIG (*n*=3, $p^{*}<0.05$, $p^{**}<0.01$, $p^{***}<0.001$).

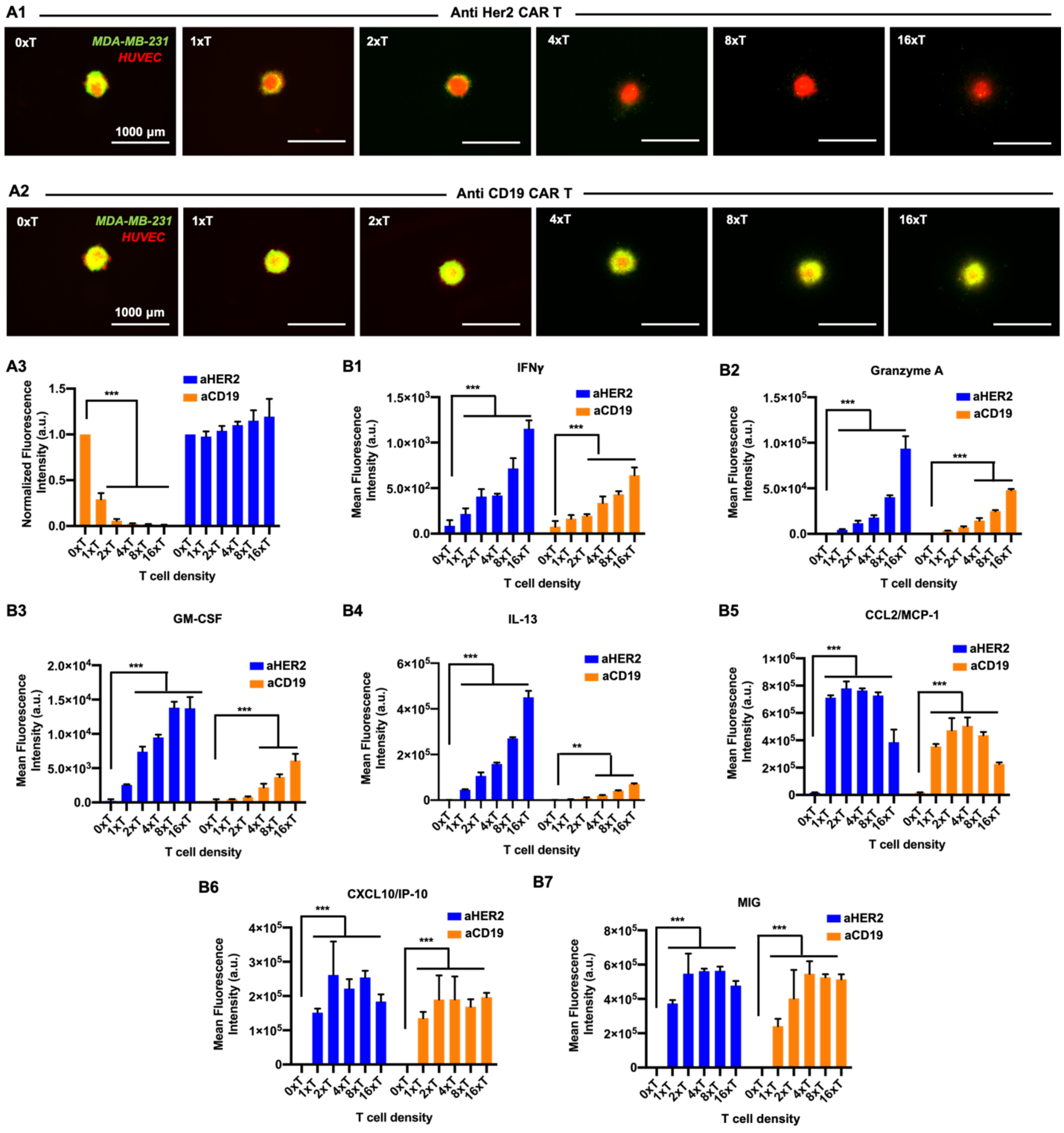


**Figure S7**: Adoptive T-cell therapy on *in vitro* static cultures of heterotypic H231F tumors. CAR-T cell density was varied as 1xT (1:1), 2xT (2:1), 4xT (4:1), 8xT (8:1), 16xT (16:1). 0xT denotes the non-treated control culture. Fluorescent images of H231F tumors after 72 h of (A1) anti HER2 CAR-T cell and (A2) anti CD19 CAR-T cell treatment. (A3) Graphical representation of normalized fluorescence intensity of anti HER2 CAR-T cell and anti CD19 CAR-T cell treatment. Graphical representation of the mean fluorescence intensity of the cytokines and chemokines secreted after 72 h of CAR-T cell perfusion including (B1) IFNγ, (B2) Granzyme A, (B3) GM-CSF, (B4) IL-13, (B5) CCL2/ MCP-1, (B6) CXCL10/ IP-10, and (B7) MIG ($n$=3, $p^{*}<0.05$, $p^{**}<0.01$, $p^{***}<0.001$).

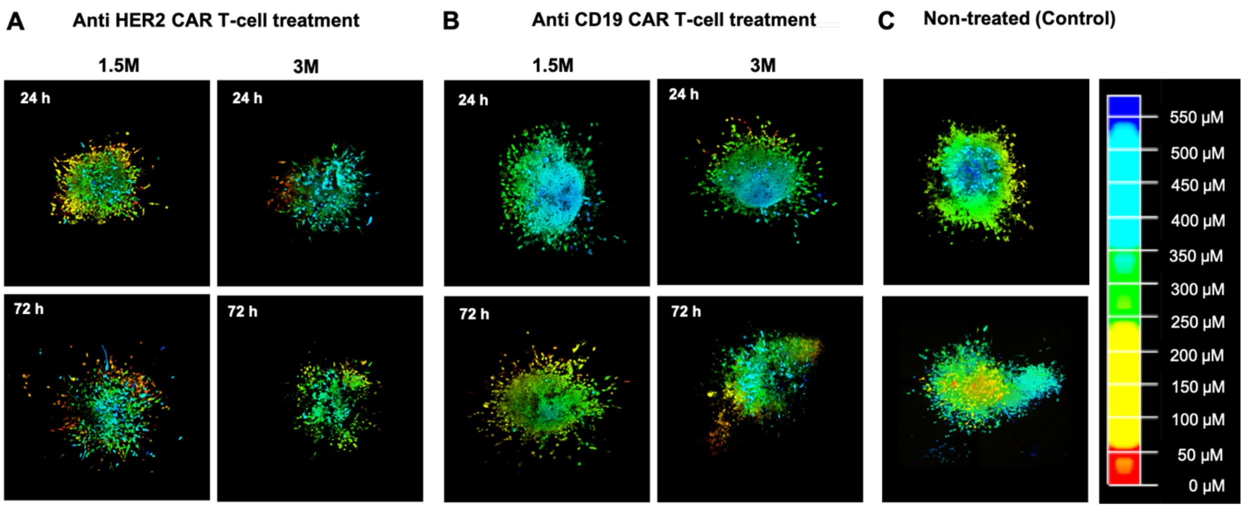


**Figure S8**: 3D Reconstruction of the entire tumor remaining after 24 or 72 h of (A) anti HER2 CAR-T cell and (B) anti CD19 CAR-T cell treatment and subsequent (C) non-treated (control) tumors. Tumors were depth coded to represent their entire volume in 3D.

### Supplementary Movies

**Movie S1**. Aspiration-assisted bioprinting of H231F tumors in a biomimetic C2F3 matrix.

**Movie S2**. 3D Reconstruction of the bioprinted perfusable tumor model illustrating the endothelialized vasculature and tumors (with angiogenic sprouting) bioprinted near the vasculature.

**Movie S3**. 3D Reconstruction of the endothelialized vasculature illustrating cancer cells trying to invade into the perfused vasculature.

**Movie S4**. CAR-T cells flowing through the endothelialized vasculature.

### References

1. N. Rajan, J. Habermehl, M.-F. Coté, C. J. Doillon, D. Mantovani, Preparation of ready-to-use, storable and reconstituted type I collagen from rat tail tendon for tissue engineering applications. *Nat. Protoc.* **1**, 2753–8 (2006).
